# Parasite hybridization promotes spreading of endosymbiotic viruses

**DOI:** 10.1101/2023.03.24.534103

**Authors:** Senne Heeren, Ilse Maes, Mandy Sanders, Lon-Fye Lye, Jorge Arevalo, Alejandro Llanos-Cuentas, Lineth Garcia, Philippe Lemey, Stephen M Beverley, James A Cotton, Jean-Claude Dujardin, Frederik Van den Broeck

## Abstract

Viruses are the most abundant biological entities on Earth and play a significant role in the evolution of many organisms and ecosystems. In pathogenic protozoa, the presence of endosymbiotic viruses has been linked to an increased risk of treatment failure and severe clinical outcome. Here, we studied the molecular epidemiology of the zoonotic disease cutaneous leishmaniasis in Peru and Bolivia through a joint evolutionary analysis of *Leishmania braziliensis* parasites and their endosymbiotic *Leishmania* RNA virus. We show that parasite populations circulate in isolated pockets of suitable habitat and are associated with single viral lineages that appear in low prevalence. In contrast, groups of hybrid parasites were geographically and ecologically dispersed, and commonly infected from a pool of genetically diverse viruses. Our results suggest that parasite hybridization, likely due to increased human migration and ecological perturbations, increased the frequency of endosymbiotic interactions known to play a key role in disease severity.

## INTRODUCTION

Viruses have the ability to infect virtually any cellular life form on Earth. Particularly fascinating are RNA viruses that infect simple eukaryotes ^1–3^, some of which have important biological roles such as limiting fungal pathogenicity ^4^ or increasing protist fecundity ^5^. Among the RNA viruses, the double-stranded RNA (dsRNA) totivirids have evolved significant diversity and are present in phyla separated by billion years of evolution, with closely related viruses identified in almost all genera of yeasts, fungi and protozoa studied ^6^. Totivirids have no lytic infectious phase and thus adopted a lifestyle of coexistence favoring long-term symbiotic persistence, passing from cell to cell mainly through mating and cell division ^7^. Because of this intimate association, it is postulated that these viruses have a mutualistic co-evolutionary history with their hosts ^8^.

Totivirids encompass most viral endosymbionts identified in pathogenic protozoa causing widespread severe illnesses such as trichomoniasis, giardiasis and leishmaniasis ^6^. An icon group of human pathogenic parasites is the genus *Leishmania* (Family Trypanosomatidae), causing the vector-borne disease leishmaniasis in about 88 countries, mainly in the tropics and subtropics ^9^. Members of the *Leishmania* genus are associated with the *Leishmania* RNA virus (LRV) (Family *Totiviridae*), forming a tripartite symbiosis with the mammalian or arthropod host. Phylogenetic studies suggest that the virus was most likely present in the common ancestor of *Leishmania*, prior to the divergence of these parasites into different species around the world ^8, 10^. This is reflected by the two types of viruses (LRV1 and LRV2) that are carried by members of the subgenera *Viannia* and *Leishmania*, respectively ^11^. It was shown that the dsRNA of LRV1 is recognized by Toll-like receptor 3 (TLR3) which directly activates a hyperinflammatory response causing increased disease pathology, parasite numbers and immune response in murine models ^12^. In human infections, the presence of LRV1 has been associated with an increased risk of drug-treatment failures and acute pathology ^13, 14^. The virus thus confers survival advantage to *Leishmania* and plays a key role in the severity of human leishmaniasis ^15, 16^.

Given the epidemiological and biomedical relevance of the *Leishmania*-LRV symbiosis ^17^, there is a clear need to understand the diversity and dissemination of the virus in parasite populations. To this end, we investigated the joint evolutionary history of *L. braziliensis* (*Lb*) parasites and LRV1 using whole genome sequencing data. The *Lb* parasite is a zoonotic pathogen circulating mainly in rodents and other wild mammals (e.g. marsupials) in Neotropical rainforests ^18^. The parasite is the major cause of cutaneous leishmaniasis (CL) in Central and South America and occasionally develops the disfiguring mucocutaneous disease where the parasite spreads to mucosal tissue. Our previous work in Peru has shown that LRV1 was present in >25% of sampled *Lb* parasites, and that the virus was significantly associated with an increased risk of treatment failure ^14^. Here, we characterize the co-evolutionary dynamics between *Lb* and LRV1 from Peru and Bolivia.

## RESULTS

### Natural genome diversity of *Lb* parasites

Our study included 79 *Lb* isolates (Supp. Table 1; Supp. Fig. 1) from Peru (N=55) and Bolivia (N=24) that were sampled during various studies on the genetics and epidemiology of leishmaniasis ^14, 19–23^, and for which the LRV infection status was previously characterized ^14^. Sequence reads of *Lb* were aligned against the *Lb* M2904 reference genome, resulting in a median coverage of 58x (min=35x, max=121x).

Variant discovery was done with GATK HaplotypeCaller to uncover high-quality Single Nucleotide Polymorphisms (SNPs) and small insertions/deletions (INDELs). The number of SNPs identified between each genome and the M2904 reference genome was relatively consistent across the panel, varying between 97,777 SNPs in CEN002 and 109,798 SNPs in LC2319 (median = 105,096; mean = 106,530). Exceptions were isolates PER231 (126,178), LC2318 (128,544) and CUM68 (134,882) that showed larger SNP densities and double the number of heterozygous sites compared to the other isolates (Supp. Fig. 2). When investigating the genome-wide distribution of normalized allele frequencies at heterozygous sites (which should be centered around 0.5 in diploid individuals) ^24^, we discovered that isolates CUM68, LC2318 and PER231 showed unbalanced read counts, symptomatic of tetraploidy ^24^, although we could not rule out contamination or a mixed infection (Supp. Fig. 3). Removing these three isolates for downstream analyses, the resulting dataset comprised a total of 407,070 SNPs and 69,604 bi-allelic INDELs called across 76 *Lb* genomes. The SNP allele frequency spectrum was dominated by low-frequency variants, with over 66% of SNPs being at <= 1% MAF. Respectively 41.5% and 5.7% of the SNPs and INDELs were found in the coding region of the genome, including 437 SNPs and 55 INDELs with a deleterious impact (e.g. introducing stop codons). Most of these deleterious mutations were rare in our panel of parasites (83.6% being at <= 1% MAF), and the remaining 80 mutations were not linked to *Lb* population structure or LRV infection status (Supp. Table 2).

Chromosome and gene copy numbers were investigated using normalized median read depths. This revealed that the majority of chromosomes were diploid (Supp. Fig. 4). Chromosome 31 was highly polysomic with at least four copies present in all isolates, a consistent observation in *Leishmania* ^25, 26^. Chromosomes 10, 26, 32 and 34 were diploid in all *Lb* isolates (Supp. Fig. 4). Excluding chromosome 31, we found 21 *Lb* isolates that were entirely diploid and 25 *Lb* isolates with at least 5 polysomic chromosomes, including 6 *Lb* isolates with at least 10 polysomic chromosomes (Supp. Fig. 4). When investigating variation in copy numbers for 8,573 coding DNA sequences, we found 201-286 amplifications and 13-33 deletions per *Lb* isolate (Supp. Table. 3), as expected for *Leishmania* and other eukaryotic genomes ^24^. Variation in chromosome and gene copy numbers was not associated with *Lb* population structure or LRV infection status.

### Epidemic clones against a background of prevalent recombination in *Lb*

It has been postulated since 1990 that protozoan parasites may have a predominantly clonal mode of reproduction and that sexual recombination events are rare ^27^, although this theory has been the subject of intense debate for more than 30 years ^28^. For *Lb,* studies using multilocus microsatellite profiles revealed contradictory results, including moderate degrees of inbreeding in populations from Peru and Bolivia ^19, 29^ and significant levels of recombination in populations from the Brazilian Atlantic Coast ^30^. These results may be biased due to the presence of population substructure (Wahlund effect) or because of the low discriminatory power of the molecular markers used ^28^.

Our phylogenetic network based on genome-wide SNPs revealed a star-like topology whereby the majority of isolates were separated by long branches (Fig. 1A), a pattern symptomatic of recombination. Indeed, levels of linkage-disequilibrium (LD) were low (*r*^2^ decayed to <0.1 within <20bp) (Supp. Fig. 5) and distributions of per-site inbreeding coefficients per population were unimodal and centered around zero (Supp. Fig. 6; Supp. Table 4), after correcting for population structure and spatio-temporal Wahlund effects. Our genome-scale data thus indicate that the distinct populations in Peru and Bolivia are approximately in Hardy-Weinberg and linkage equilibrium, suggesting that recombination may be a prevalent and universal process in this species.

**Figure 1.**
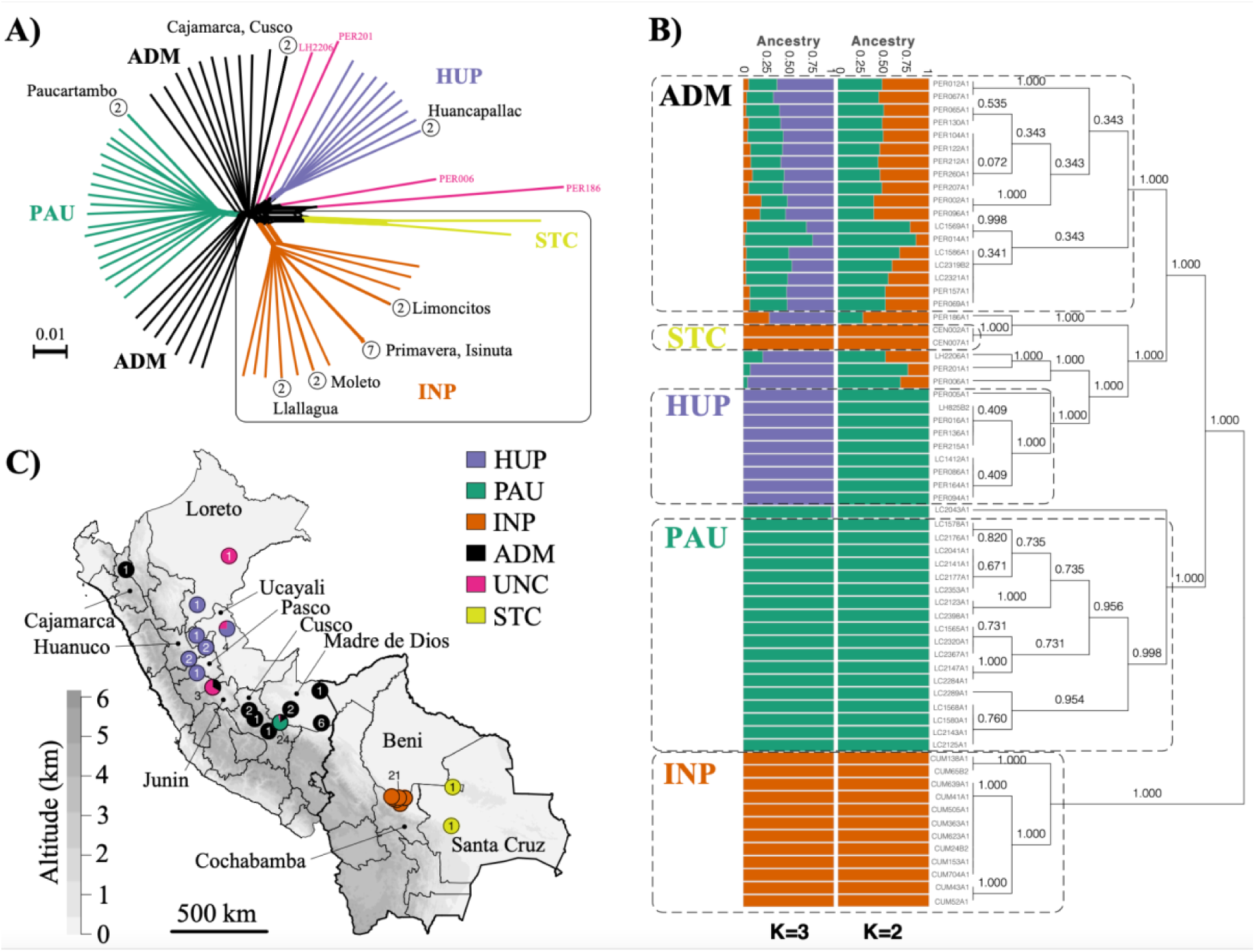
Population genomic structure and admixture in *Lb* parasites from Peru and Bolivia. **A)** Phylogenetic network as inferred with SPLITSTREE based on uncorrected *p*-distances between 76 *Lb* genomes (excluding isolates CUM68, LC2318 and PER231) typed at 407,070 bi-allelic SNPs. Branches are coloured according to groups of *Lb* parasites as inferred with ADMIXTURE and fineSTRUCTURE (panel B). Box indicates the position of the Bolivian *Lb* genomes; all other genomes were sampled in Peru. Circles at the tips of seven branches point to groups of near-identical *Lb* genomes, showing the number of isolates and their geographical origin. **B)** ADMIXTURE barplot summarizing the ancestry components inferred at *K* = 2 or *K* = 3 populations in 65 *Lb* genomes. The phylogenetic tree summarizes the fineSTRUCTURE clustering results based on the haplotype co-ancestry matrix. Numbers indicate the MCMC posterior probability of a given clade. Dashed boxes indicate the three main parasite groups (INP, HUP, PAU) and the two main groups of admixed parasites (ADM, STC); the remainder of the parasites were of uncertain ancestry (UNC). **C)** Geographic map of Peru and Bolivia showing the origin of 76 *Lb* genomes. Dots are coloured according to groups of *Lb* parasites as inferred with ADMIXTURE and fineSTRUCTURE (panel B). Gray-scale represents altitude in kilometers, indicating the position of the Andes along the Peruvian and Bolivian Coast. Names are given for those Peruvian departments where a *Lb* parasite was isolated.

While the majority of *Lb* genomes differed at an average 9,866 fixed SNP differences (median 9,494), we also identified seven small groups of isolates with near-identical genomes that cluster terminally in the phylogenetic network (Fig. 1A, circles with numbers). Isolates within each of these groups displayed no or few fixed SNP differences and a relatively small amount of heterozygous SNP differences (Supp. Table 5), and are thus likely the result of clonal propagation in the wild. With the exception of a pair of near-identical genomes found in two distant Departments of Peru (Cajamarca and Cusco), each of the clonal lineages were geographically confined (Fig. 1A; Supp. Table 1). This is best exemplified by four clonal lineages that each circulated in a different location in the National Park of Isiboro in Bolivia (Fig. 1A). One of these lineages was sampled over a period of 12 years during an epidemic outbreak of leishmaniasis in Cochabamba (Bolivia), suggesting that the observed clonal population structure is temporally stable (Supp. Table 5).

Our data thus suggest that *Lb* population structure follows an epidemic/semi-clonal model of evolution ^31, 32^, as proposed for other protozoan parasites ^33^. This model assumes frequent recombination within all members of a given population, but that occasionally a successful individual increases in frequency to produce an epidemic clone ^31, 32^.

### Parasite populations are isolated in pockets of suitable habitat

Divergent *Lb* ecotypes were described in Peru ^20, 21^ and Eastern Brazil ^34^ that are each associated with particular ecological niches, suggesting that the environment may play a key role in *Lb* population structure. In Peru, our previous work revealed the existence of two Andean and one Amazonian *Lb* lineage ^20^. Here, we characterized in more detail the ancestry of the Amazonian *Lb* parasites.

We included one isolate from each clonal lineage and removed SNPs showing strong Linkage Disequilibrium (LD), leaving a total of 176,143 SNPs for investigating population structure. We identify three distinct ancestry components (PAU, HUP, INP) corresponding to groups of *Lb* parasites with geographically restricted distributions (Fig. 1B,C; Supp. Fig 7). These three populations showed signatures of spatial and temporal genetic structure: PAU (N=19) was sampled between 1991 and 1994 in Paucartambo (Southern Peru), INP (N=21) was sampled between 1994 and 2002 in the Isiboro National Park (Central Bolivia) and HUP (N=10) was sampled between 1990 and 2003 in Huanuco, Ucayali and Pasco (Central Peru) (Fig. 1B,C; Supp. Fig 7). Assuming *K* = 2 populations, the two Peruvian populations PAU and HUP were grouped as one population (Fig. 1B). While our results indicate that *Lb* may be genetically clustered at sampling sites, we also identify several groups of *Lb* parasites with uncertain ancestries (ADM, UNC and STC), one of which contained a total of 19 *Lb* isolates sampled between 1991 and 2003 across Central and Southern Peru (ADM) (Fig. 1B,C).

The strong signatures of geographical isolation of the three inferred ancestry components (PAU, INP and HUP) prompted us to investigate the geographical and/or environmental variables that impacted the population structure of *Lb* in the region. This was done through redundancy analysis (RDA) and generalized dissimilarity modeling (GDM), including geographic distance and 19 bioclimatic variables (Suppl Table 6). Variable selection using two approaches consistently pinpointed differences in isothermality (bio3) and precipitation of driest month (bio14) between sampling locations as the environmental contributors to parasite genetic distance (Suppl Table 7; Suppl. Results). The RDA model revealed that environmental differences and geographic distance explained one-third (27.3%) of the genomic variability, of which 10.2% was contributed by environmental differences and 7.5% by geographic distance (Suppl Fig 8A; Suppl. Table 8; Suppl. Results). Similar results were obtained with the GDM model (Suppl. Results).

Environmental niche modeling (ENM) using present and past bioclimatic variables predicted the putative suitable habitat of the *Lb* populations during the Last Glacial Maximum (LGM; 21 kya) and the LIG (130 kya). Present-day predicted suitability regions for *Lb* coincided with tropical rainforests as predicted by the Köppen-Geiger (KG) climate classification, and shows that the three *Lb* populations are surrounded by the less suitable tropical monsoon forests and the non-suitable Andean ecoregions (Fig 2; Supp. Fig. 9, 10). Suitability predictions for the LGM and the LIG periods revealed largely similar ecological niches, suggesting that *Lb* suitable habitats have been relatively stable over the past 130,000 years (Supp. Fig. 9, 10).

**Figure 2.**
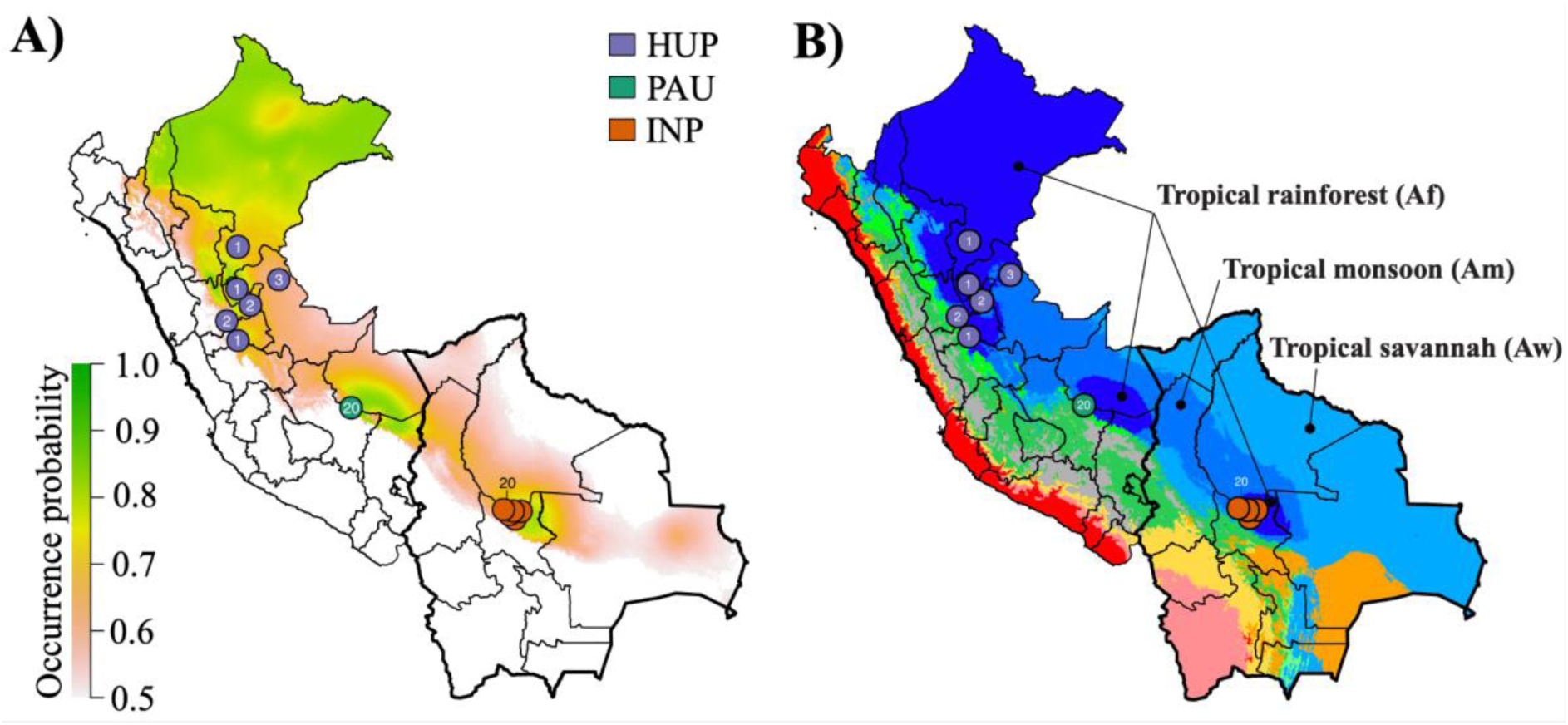
Distribution of *Lb* parasite populations and their endosymbiotic LRV1 lineages in Peru and Bolivia. **A)** Present-day predicted suitability regions for *Lb* parasites as revealed with ecological niche modeling. The continuous-scale legend represents habitat suitability (probability of occurrence). Map shows the distribution of isolates belonging to the three *Lb* parasite populations HUP, PAU and INP. **B)** Ecological regions in Peru and Bolivia as per Köppen-Geiger climate classification. Only three ecological regions were labeled for visibility reasons. The distribution is shown for *Lb* isolates belonging to the three *Lb* parasite populations HUP, PAU and INP.

### Hybrid parasites are geographically and ecologically dispersed

In addition to the three *Lb* populations, we also identified three groups of *Lb* parasites with mixed ancestries: one large group of 19 *Lb* isolates (ADM) sampled between 1991 and 2003 mainly from Southern Peru (Junin, Cusco and Madre de Dios), four *Lb* isolates (UNC) sampled between 2001 and 2003 from Central/Northern Peru (Junin, Ucayali and Loreto) and one group of two isolates (STC) sampled in 1984/1985 from the Santa Cruz Department in Central Bolivia (Fig. 1). The genetic diversity of the largest group of hybrid parasites (ADM) was much larger compared to the three *Lb* populations: the number of mitochondrial haplotypes and nuclear segregating sites was 1.4-4 times higher in the ADM group (309,543 SNPs; 8 haplotypes) compared to that of the PAU (189.748 SNPs; 4 haplotypes), INP (185.927 SNPs; 4 haplotypes) and HUP (215.741 SNPs; 2 haplotypes) populations. The observation of a large number of mitochondrial haplotypes in the ADM group suggests that these parasites are descendants from multiple independent hybridization events (Supp. Fig. 11).

We used PCAdmix to infer the genome-wide ancestry of all admixed individuals and three control individuals from each source (Fig. 3). Ancestry was assigned to phased haplotype blocks of 30 SNPs by comparing them to reference panels of PAU, INP and HUP. While the control samples were assigned 92.9-99.4% to their respective populations, the admixed individuals showed mixed ancestries between PAU (21.1-73.1%), INP (14.3-36.7%) and HUP (0.1-54.4%). Plots of local ancestry revealed a complex and heterogeneous pattern of mosaic ancestry between the three sources (Fig. 3B, Supp. Fig. 12), suggesting that the hybrid parasites experienced many cycles of recombination following the initial admixture event(s). We next used the three-population statistic *f*_3_ to formally test the potential source populations for introgressed alleles in the ADM and STC groups. When testing (test; A,B), a negative result indicates that the test group is an admixed population from A and B. We found a significantly negative *f*_3_ statistic when ADM was the test group with PAU/HUP (*f*_3_ = -0.0013, Z-score = -18.2) and PAU/INP (*f*_3_ = -0.0002, Z-score = -2.8) as sources, but not with INP/HUP (*f*_3_ = 0.0013, SD=, Z-score = 14.8) as sources. All estimated *f*_3_ statistics were positive when STC was used as the test group. Hence, the ancestry of STC remains largely unresolved, but may involve admixture with divergent *Lb* parasites not captured in this study, as indicated by their distant position in a haplotype network of mitochondrial maxicircle SNPs (Supp. Fig. 11) and by the comparatively large number of fixed SNP differences between STC and other *Lb* groups (Supp. Fig. 13).

**Figure 3.**
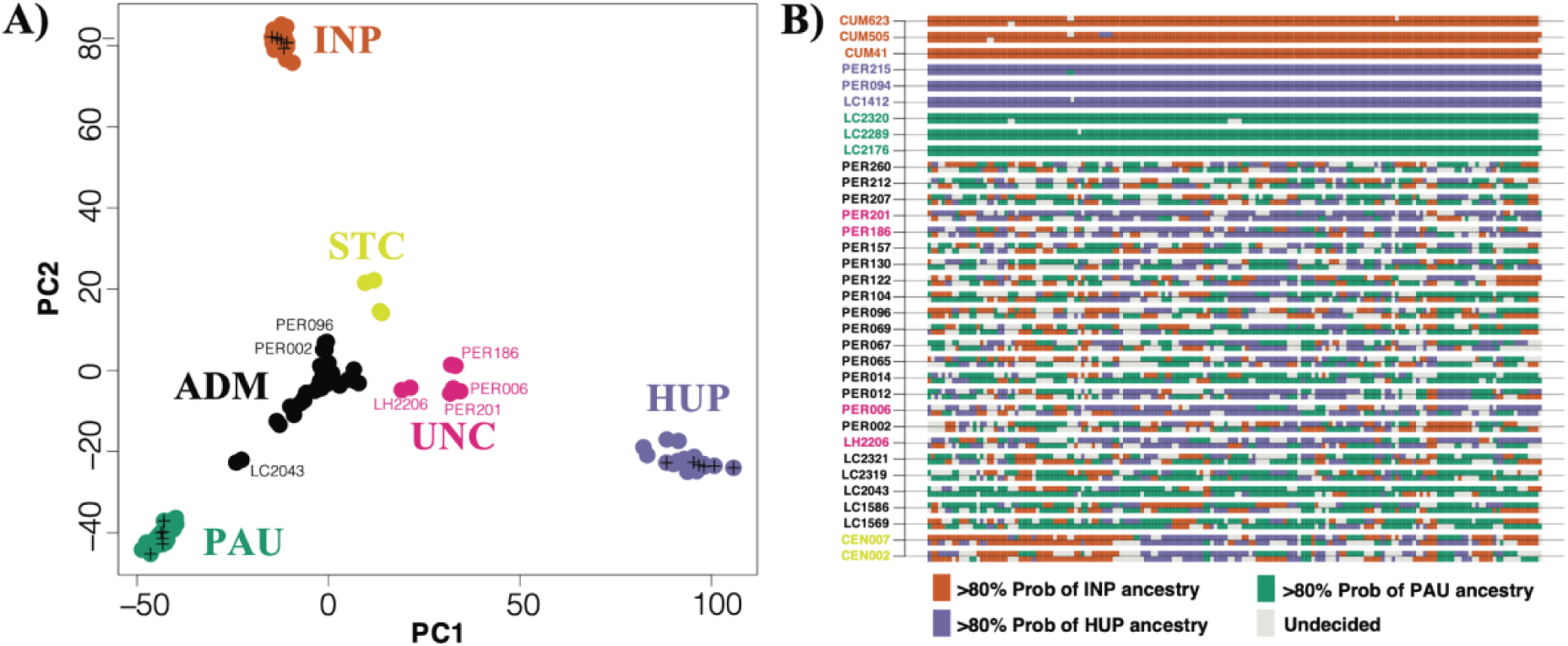
Mosaic ancestry in hybrid *Lb* parasites. **A)** PCA-based ancestry assignment of parasites of uncertain ancestry (ADM, UNC, STC) assuming three *Lb* populations (ADM, INP, HUP). Plus signs indicate three randomly selected genomes of each source population that were used as controls. **B)** PCA-based local ancestry assignment to PAU, INP and HUP source populations of the 23 ADM isolates, 2 STC isolates and three randomly selected isolates from each source population. Ancestry was assigned in windows of 30 SNPs along chromosome 35 (examples for other chromosomes are shown in Supp. Fig. 12).

The distribution of the three groups of uncertain ancestries largely pair with the distribution of the *Lb* populations: the ADM group is mainly found within ecological regions surrounding the location of the PAU population (Southern Peru), the UNC group is found within ecological regions surrounding the location of the HUP population (Central/Northern Peru) and the STC group is found near the INP population (Central Bolivia) (Fig. 1C). In addition, the median distance between hybrids from the ADM group (310 km), STC group (203.8 km) and UNC group (349 km) was much larger compared to the median distance between parasites of each of the populations PAU (0 km), HUP (155 km) and INP (37 km).

Altogether, our observations show that hybrid parasites are geographically and ecologically more widespread compared to the three parasite populations, suggesting that secondary contacts occurred following migration out of the suitable rainforest habitats.

### High viral prevalence and lineage diversity within group of hybrid parasites

A total of 31 out of 79 included *Lb* isolates (39%) were LRV1+, as revealed by previous work ^14^. While these originated mainly from Peru (N=27), the geographic distribution of LRV1+ and LRV1-*Lb* isolates was approximately similar (Supp. Fig. 1). Here, we recovered LRV1 genomes for 29/31 LRV1+ *Lb* isolates from Peru and Bolivia following a *de novo* assembly of total RNA sequencing reads (Supp. Table 9; Supp. Results). The procedure failed for two *Lb* isolates, either because of difficulties in growing cultures (PER096) or because assembly yielded a partial LRV1 genome (PER231). Two *Lb* isolates (CUM65 and LC2321) each harbored two LRV1 genomes (Supp. Results), differing at 999 (for CUM65) and 60 (for LC2321) nucleotides, bringing the total to 31 viral genomes. While only 0.04% (0.004%-0.1%) of the RNA sequencing reads aligned against the LRV1 assemblies, median coverages were relatively high, ranging between 31X and 868X (median = 372X) (Supp. Table 9). Analogous genomic regions in the assembled sequences were identical to ∼1 kb sequences as obtained with conventional Sanger sequencing, confirming the high quality of our assemblies (Supp. Results).

Sequences were 4,738 - 5,285 bp long, covering the full-length coding sequence of the virus, and showing an average GC content of 46% (45.4%-46.8%) (Supp. Table 9). All but two genomes included two open reading frames without internal stop codons, encoding the Capsid Protein (CP; 2,229bp) and the RNA-dependent RNA Polymerase (RDRP; 2,637bp). Two isolates (PER014 and PER201) each contained 1 internal stop codon at amino acid positions 875 (TAA) and 882 (TAG), respectively. Sequence identities between these novel LRV1 genomes from Peru and Bolivia, and a previously published LRV1 genome from French Guiana (YA70; KY750610) ranged between 80% and 81%. Amino acid identity of both genes against YA70 ranged between 94%-96% for CP, and 85%-87% for RDRP (Supp. Table 9).

We defined a total of nine divergent viral lineages in Peru and Bolivia (L1-9), all supported by high bootstrap values (Fig. 4) and low pairwise genetic distances (<0.09; Supp. Table 10; Supp. Table 11). No evidence of recombination was found in our set of LRV1 sequences, following pairwise homoplasy index (PHI) tests (p=0.99). The number of LRV1 lineages sampled per locality was associated with the number of sampled parasites (t=5.72; p < 0.001) (Supp. Figure 14). For instance, the most densely sampled location (Paucartambo, Cusco, Peru) in terms of *Lb* parasites (N = 25) also contained the most viral lineages (Supp. Figure 14), two of which (L5 and L9) were only found in this location (Fig. 4). Other locations that include multiple viral lineages are Tambopata (Madre de Dios, Peru) (L3 and L4) and Moleto (Cochabamba, Bolivia) (L7, L8) (Fig. 4; Supp. Figure 14). These results show that multiple divergent viral lineages could co-occur within the same geographic region, and that a more dense sampling of parasites may uncover more viral lineages.

**Figure 4.**
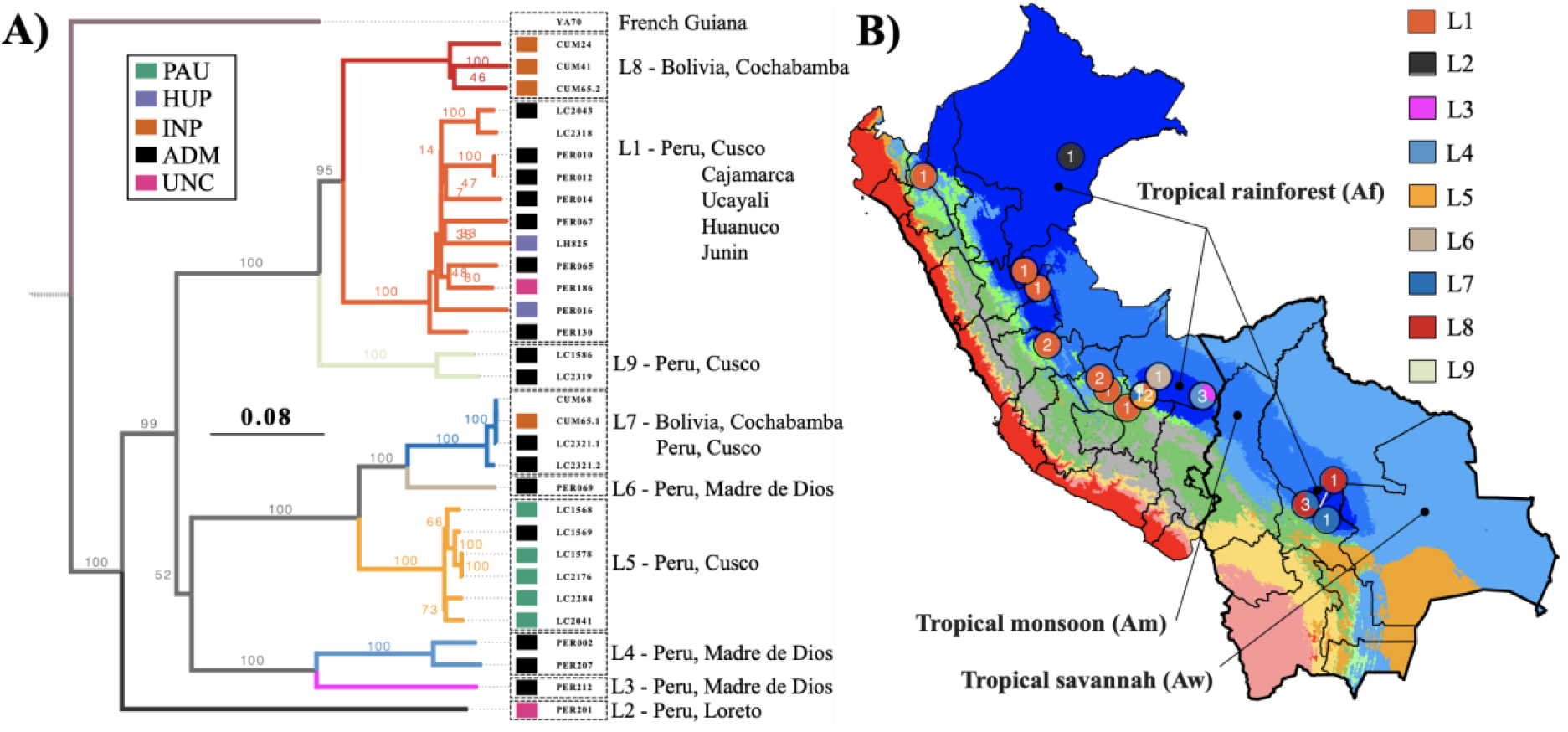
Evolutionary history of LRV1 in Peru and Bolivia. **A)** Midpoint-rooted maximum likelihood phylogenetic tree based on 31 LRV1 genomes of *Lb* from Peru and Bolivia, one LRV1 genome (YA70) of *Lb* from French Guiana and 26 LRV1 genomes of *Leishmania guyanensis* from Brazil, Suriname and French Guiana (the latter were omitted for visibility reasons). The position of the root is indicated with a dashed line. Clades are colored according to the different viral lineages (L1-L9). Colored boxes at the tips of each branch represent the population structure of *Lb* (see legend), with colors matching the different groups of parasites as shown in Figure 1. Note that the tetraploid hybrid parasites LC2318 and CUM68 from Peru and Bolivia, and isolate YA70 from French Guiana were omitted from the analyses of parasite population structure; these were thus not assigned to any parasite group. **B)** Ecological regions in Peru and Bolivia as per Köppen-Geiger climate classification. Only three ecological regions were labeled for visibility reasons. The distribution is shown for LRV+ *Lb* isolates belonging to the nine viral lineages L1-L9.

The majority of viral lineages (L2-6, L8-9) were found in a single locality (Fig. 4), suggesting that the geographic spread of most LRV1 lineages is restricted. Two viral lineages were more widely distributed: one large group of viral strains (L1) was found along the Andes from Northern Peru to Southern Peru, and one group (L7) was found in Cusco (Peru) and Cochabamba (Bolivia) (Fig. 4). Three viral strains of lineage L7 were virtually identical: viral sequences of the Bolivian *Lb* parasites CUM68 and CUM65 were identical, and differed by one nucleotide from a viral sequence of the Peruvian *Lb* strain LC2321. The distal position of the two Bolivian L7 strains within a clade of Peruvian viral lineages suggests that the L7 viral lineage was introduced in Bolivia from Southern Peru (Fig. 4). Finally, all early-diverging lineages (L2-L9) were found in the tropical rainforests (KG-Af) of Peru and Bolivia, except for the widespread lineage L1 that was found in tropical rainforests (KG-Af), tropical monsoon (KG-Am), tropical savannah (KG-Aw) and temperate climate (KG-Cwb) (Fig. 4). This suggests that LRV1 predominantly evolved in the lowland tropical rainforests before spreading to other ecological regions.

When investigating the distribution of viral lineages across the different *Lb* parasite groups, we found two impactful observations. First, LRV1 prevalence differs between the different *Lb* groups, with a significantly lower prevalence in the *Lb* populations PAU (30%; 6/20), INP (14.3%; 3/21) and HUP (20%; 2/10) compared to the ADM (78.9%; 15/19) and UNC (50%; 2/4) groups that included *Lb* parasites of uncertain ancestry. Second, the three *Lb* populations PAU, INP and HUP (comprising a total of 50 *Lb* isolates) each consisted of strictly one viral lineage: the two LRV+ *Lb* isolates from HUP comprised the L1 viral lineage, the five LRV+ *Lb* isolates from PAU comprised the L5 viral lineage and the three LRV+ *Lb* isolates from INP comprised the L8 viral lineage, with the exception of isolate CUM65 that was coinfected with a viral strain from lineage L7 (Fig. 4). These observations strongly suggest that the three isolated *Lb* populations are each associated with a unique viral lineage. In contrast, the group of widespread hybrid *Lb* parasites (ADM) comprised almost all viral lineages (L1, L3-L9), four of which were found exclusively in the ADM *Lb* group (Fig. 4). The extensive viral lineage diversity within the ADM group suggests that parasite gene flow and hybridization has promoted the dissemination of viral lineages. This is best exemplified by the widespread L1 viral lineage that is mainly associated with *Lb* parasites from the ADM group, suggesting that the spread of L1 viruses along the Andes was mediated by parasite hybridization.

### Leishmaniaviruses co-diverge with Leishmania (Viannia) host species

Phylogenetic studies indicate that LRV co-diverged with *Leishmania* spp. ^35^, though the virus is not identified in all *Leishmania* species ^11, 36^ and the presence of viruses in some *Leishmania* species may be due to relatively recent horizontal transfer ^37–39^. The latter observation corroborates experimental evidence that LRV transfer among *Leishmania* species may occur via exosomes that are shed by infected *Leishmania* cells ^40^. The degree of LRV1 co-divergence between *Leishmania Viannia* species has not yet been documented.

Viral evolutionary analyses were done using partial (N = 70) and full-length genome (N = 57) sequences, including publicly available LRV1 genomes of *Lb*, *L. guyanensis* (*Lg*) and *L. shawi* (*Ls*) from Brazil, French Guiana and Suriname. Despite the allopatric sample, a phylogeny based on LRV1 genomes revealed two groups of viral strains: one including *Lb* viral strains from Peru, Bolivia and French Guiana, and one including *Lg* viral strains from Brazil, French Guiana and Suriname (Fig. 5A). Similar results were obtained based on partial sequences of LRV1, whereby viral strains from Brazil clustered either with the *Lb* or *Lg* viral strains (Supp. Fig. 15). In addition, nucleotide differences were on average higher between *Lb* and *Lg* viral genomes (median = 1,114, min = 1,065, max = 1,208) than between viral genomes of *Lb* (median = 908, min = 0, max = 991) and *Lg* (median = 647, min = 24, max = 1,156) (Supp. Fix. 16; Supp. Table 12). These results show that LRV1 consists of divergent viral lineages that are grouped by *Leishmania* host species.

**Figure 5.**
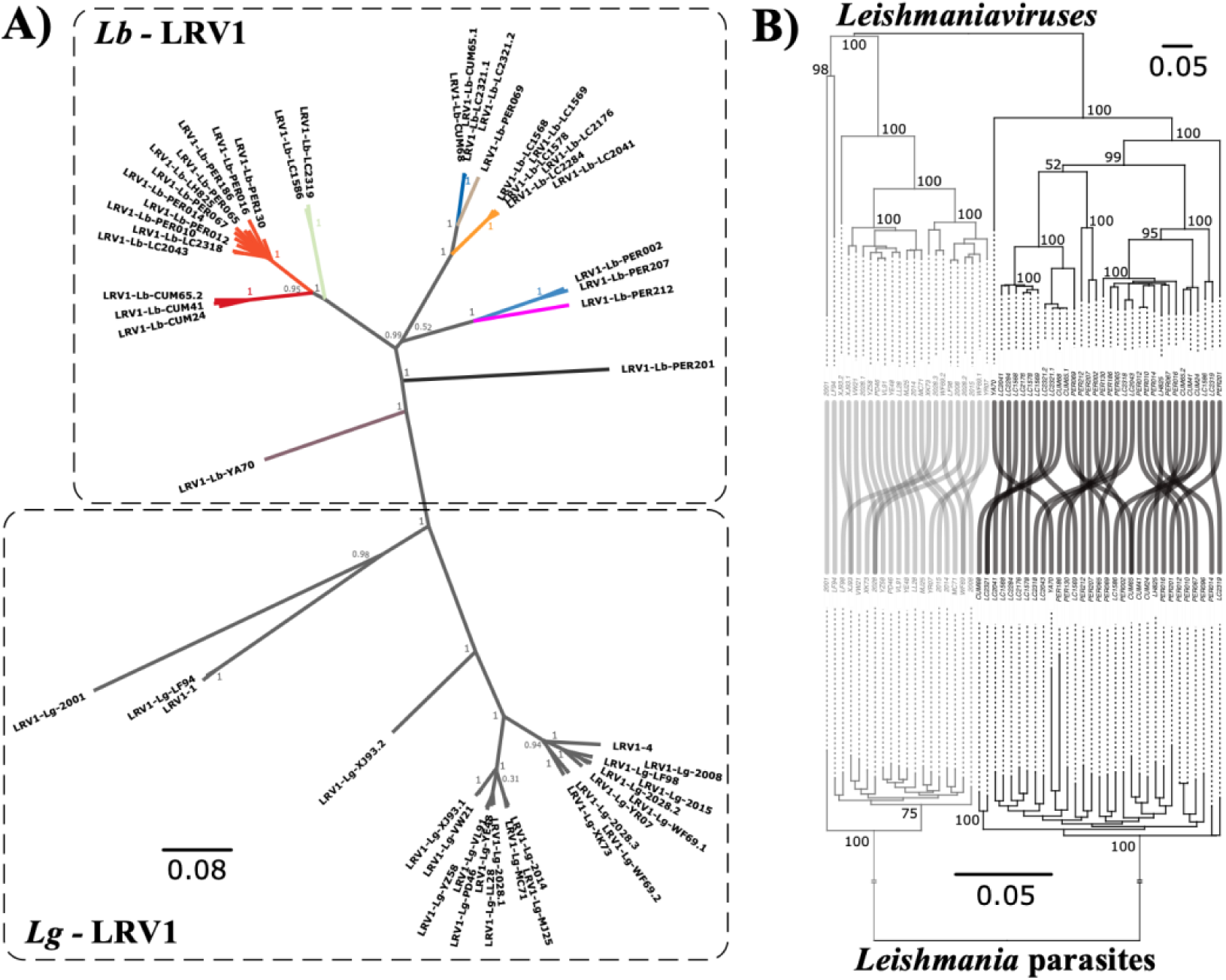
Co-divergence between *Leishmania* and LRV1. **A)** Unrooted maximum likelihood phylogenetic tree based on 31 LRV1 genomes of *Lb* from Peru and Bolivia, one LRV1 genome (YA70) of *Lb* from French Guiana and 26 LRV1 genomes of *Lg* from Brazil, Suriname and French Guiana. LRV1 clades of *Lb* are coloured according to the different viral lineages identified in this study (L1-L9) (see Figure 4). **B)** Phylogenetic tangle plot revealing clear patterns of virus-parasite co-divergence between *Lb* and *Lg* and their respective monophyletic LRV1 clades. The gray and black branches and tree links correspond to *Lg* and *Lb*, respectively.

In order to investigate whether LRV1 co-diverged with *Lb* and *Lg* parasites, we reconstructed phylogenies based on *Lb*+*Lg* viral genomes and their corresponding *Lb*+*Lg* host genotypes. *Leishmania* genotypes were based on SNPs called with GATK Haplotypecaller across our set of 79 *Lb* genomes and a set of 20 publicly available *Lg* genomes (see methods). Despite the relatively low coverage of *Lg* genomes (median 3x, min 1x, max 4x), genotyping recovered a total of 7,571 high-quality bi-allelic SNPs called across 99 *Leishmania* genomes. Similar to results obtained for LRV1 (Fig. 5A), a phylogenetic network revealed a clear dichotomy between *Lb* and *Lg* parasites (Supp. Fig. 17). Co-phylogenetic analyses revealed a split of both viral and parasite genomes at the deepest phylogenetic node, confirming that LRV1 viruses cluster with their *Leishmania* host species (Fig. 5B). A consecutive permutation test for co-speciation confirmed the topological similarity between both phylogenetic trees (RF=70, p-value = 0.001). The extremely low coverage of publicly available sequence data for *Lg* precluded us to study intraspecific patterns of co-divergence and host-switching within a co-phylogenetic framework.

## DISCUSSION

Our main goal was to understand the evolution and dissemination of endosymbiotic viruses (here LRV1) in an important group of human pathogenic parasites (here *Lb*). Genetic studies have shown that both *Lb* ^20, 29, 30, 34, 41^ and LRV1 ^43^ populations are structured according to their geographical origin. Here, we characterized the joint ancestry of the two species in Peru and Bolivia based on whole genome sequence analysis. Our results show that both *Lb* and LRV1 are genetically highly heterogeneous and predominantly evolved within the lowland tropical rainforests. Models of landscape genomics reveal that geographic distance and in particular environmental differences between sampling locations contributed to partitioning *Lb* diversity. This indicates that the large diversity and population substructure of *Lb* and LRV1 may have been driven by the extremely diversified ecosystem of the Amazonian rainforest, including various host–vector communities ^44^.

The degree of LRV1 co-divergence and host-switching between and within *Leishmania* species has not yet been documented. We show that LRV1 lineages cluster according to different *Leishmania* species, indicating that horizontal transfer of LRV1 between parasite species is rare ^43, 45^. The agreement between LRV1 and *Leishmania* phylogenies corroborates the general hypothesis that LRV1 is an ancient virus that has co-evolved with its parasite ^8, 11, 35^. While we observed a clear pattern of co-divergence at the species level, we did not detect such a pattern when investigating co-phylogenies of LRV1 and *Lb*. Our data suggest that recombination in *Lb* may drive prevalent horizontal transmission of LRV1 at the intraspecific level. Such intraspecific phylogenetic incongruences have also been observed for other endosymbionts, such as *Wolbachia* ^46^.

One third of our panel showed signatures of mixed ancestry, supporting a growing body of genomic evidence for extensive genetic exchange in protozoan parasites in the wild ^20, 28, 47–52^. We propose that increased human migration and ecological perturbations over the past decades, including movement of hemerophile reservoirs/hosts such as dogs and rats, may have resulted in the displacement of *Lb* parasites out of the tropical rainforests ^29, 53^. Probably the most exciting outcome of our study is that hybrid parasites were geographically and ecologically widely distributed and commonly infected from a pool of genetically diverse LRV1, in contrast to the *Lb* populations that were confined to tropical rainforests and infrequently associated with a single LRV1 lineage. Our data thus suggests that parasite hybridization increased the frequency of *Lb*-LRV1 symbiotic interactions, which play a key role in the severity of human leishmaniasis ^15, 16^ and which may have profound epidemiological consequences in the region.

## METHODS

### Nucleic acid extraction and sequencing of DNA parasites and their dsRNA viruses

This study included 79 *Lb* isolates (Supp. Table 1) from Peru (N=55) and Bolivia (N=24) that were sampled within the context of various studies on the genetics and epidemiology of leishmaniasis. In Bolivia, the majority of *Lb* isolates (N=21) were sampled between 1994 and 2002 within the context of a CL outbreak in the Isiboro National Park (Department of Cochabamba). Two *Lb* isolates were sampled between 1984 and 1985 within the Santa Cruz Department and one is of unknown origin (Supp. Fig. 1). In Peru, half of the *Lb* isolates were sampled between 1991 and 2003 in the Cusco state (N=29), mainly from the Paucartambo province (N=25). The remaining 26 *Lb* isolates originated from Madre de Dios (N=9), Ucayali (N=5), Huanuco (N=4), Junin (N=4), Loreto (N=2), Pasco (N=1) and Cajamarca (N=1) (Supp. Fig. 1).

All 79 *Lb* isolates were cultured for 3-4 days on a HOMEM medium with Fetal Bovine Serum (FBS) at the Antwerp Institute of Tropical Medicine. Parasites were subjected to a small number of passages (mean = 18 ± 5) to reduce potential culture-related biases in parasite genomic characterization ^54^. Parasite cells were centrifuged into dry pellets and their DNA was extracted using the QIAGEN QIAmp DNA Mini kit following the manufacturer’s protocol. At the Wellcome Sanger Institute, genomic DNA was sheared into 400 to 600 base pair fragments by focused ultrasonication (Covaris Inc.), and amplification-free Illumina libraries were prepared ^55^. One hundred base pair paired-end reads were generated on the HiSeq 2000, and 150 base pair paired end reads were generated on the HiSeq ×10 according to the manufacturer’s standard sequencing protocol.

Previous work has shown that 31 out of the 79 *Lb* isolates were positive for LRV1 ^14^. The majority of the LRV1-positive *Lb* isolates originated from Peru (N=27) while the remaining four isolates were sampled in the Isiboro National Park (Cochabamba, Bolivia) (Supp. Fig 1). More than half of the LRV1-positive isolates in Peru originated from Cusco (N=15) of which 11 were sampled in the Paucartambo province. LRV1 was also sampled in Madre de Dios (N=5), Junin (N=3), Cajamarca (N=1), Huanuco (N=1), Loreto (N=1) and Ucayali (N=1) (Supp. Fig 1).

The 31 LRV1-positive *Lb* isolates ^14^ were grown on a HOMEM medium with FBS for 2-3 weeks (mean passage number = 18 ± 5) at the Antwerp Institute of Tropical Medicine, ensuring high parasite yields (ca. 10^7^-10^8^ parasites/ml). Isolation of dsRNA was performed as previously described ^56^. In short, isolation involved a TRIZOL reagent (Invitrogen) RNA extraction followed by a RNase-free DNase I (NEB.) and a S1 nuclease (Sigma-Aldrich) treatment along with an additional purification step (Zymoclean). Double-stranded RNA of approximately 5.2kb. was visualized on 0.8% agarose gel (TAE buffer) stained with ethidium bromide. Extracts of dsRNA were sequenced at Genewiz (Leipzig, Germany) using the NovaSeq platform (Illumina) generating 150bp paired end reads.

### Bioinformatics and population genomics of *Lb* parasites

Sequencing reads were mapped against the *Lb* M2904 reference genome using SMALT v.0.7.4 (https://www.sanger.ac.uk/tool/smalt-0/) as previously described ^20^. The reference genome comprises 35 chromosomes (32.73 Mb) and a complete mitochondrial maxicircle sequence (27.69 kb), and is available on https://tritrypdb.org/ as LbraziliensisMHOM/BR/75/M2904_2019. Genome wide variant calling (SNPs, INDELs) was done using GATK v.4.0.1.0 ^57, 58^. More specifically, we used GATK HaplotypeCaller for generating genotype VCF files (gVCF) for each *Lb* isolate. Individual gVCF files were combined and jointly genotyped using CombineGVCFs and GenotypeGVCFs, respectively. SNPs and INDELs were separated using SelectVariants. Low quality variants were excluded using GATK VariantFiltration following GATK’s best practices^59^ and BCFtools v.1.10.2 ^60^ view and filter. Specifically, SNPs were excluded when QD < 2, FS > 60.0, SOR > 3.0, MQ < 40.0, MQRankSum < -12.5, ReadPosRankSum < -8.0, QUAL < 250, format DP < 10, format GQ < 25, or when SNPs occurred in clusters (ClusterSize=3, clusterWindowSize=10). INDELs were excluded when QD < 2, FS < 200.0, ReadPosRankSum < -20.0. The final set of SNPs and INDELs were annotated using the *Lb* M2904 annotation file with SNPEFF v4.5 ^61^. At heterozygous SNP sites, the frequencies of the alternate allele read depths ^24^ were calculated using the vcf2freq.py script (available at github.com/FreBio/mytools).

Chromosomal and gene copy number variation were estimated using normalized read depths. To this end, per-site read depths were calculated with SAMtools depth (-a option). Haploid copy numbers (HCN) were obtained for each chromosome by dividing the median chromosomal read depth by the median genome-wide read depth. Somy variation was then obtained by multiplying HCN by two (assuming diploidy). To obtain gene HCN, the median read depth per coding DNA sequence (CDS) was divided by the median genome-wide read depth. The HCN per CDS were summed up per orthologous gene group. Gene copy number variations were then defined where the z-score was lower than -1 (deletions) or larger than 1 (amplifications).

A NeighborNet tree was reconstructed based on uncorrected *p*-distances with SplitsTree v.4.17.0^62^. The population genomic structure of *Lb* was examined with ADMIXTURE v.1.3.0^63^ and fineSTRUCTURE v.4.1.1^64^. ADMIXTURE was run assuming 1 to 10 populations (*K*), performing a 5-fold cross-validation procedure, and after removing SNPs with strong LD with plink v.1.9 ^65^ (--indep-pairwise 50 10 0.3). CHROMOPAINTER analysis (as part of fineSTRUCTURE) was run to infer the genetic ancestry based on haplotype similarity. To this end, individual genotypes were computationally phased with BEAGLE v.5.2 ^66^ using default parameters, after which fineSTRUCTURE was run using 8e06 MCMC iterations with 50% as burn-in, and 2e06 maximization steps for finding the best state for building the tree ^48^. Local ancestry was assigned with PCAdmix ^67^ using phased genotype data (i.e. haplotypes) as obtained with the BEAGLE v5.4 ^66^. *F3*-statistics were calculated with Treemix v1.13 ^68^. Finally, Hardy-Weinberg equilibrium (HWE) was examined by calculating per site the inbreeding coefficient as Fis = 1 – Ho/He; with Ho representing the observed heterozygosity and He the expected heterozygosity. Decay of LD was calculated and visualized using PopLDdecay ^69^. To control for spatio-temporal Wahlund effects, we calculated Fis and LD decay using subsets of isolates that were isolated close in time (year of isolation) and space (locality), and taking into account population genomic structure (Supp. Table 4).

To investigate the spatio-environmental impact on genetic variation among the three *Lb* populations we included geographic distances among sampling locations and extracted 19 bioclimatic variables of the WorldClim2 database ^70^. We firstly investigated the role of geography on the genomic differentiation of the *Lb* components through linear and non-linear regression analyses of distance matrices (Supp. Methods). Secondly, we used redundancy analysis (RDA) ^71^ and generalized dissimilarity modeling (GDM) ^72^ to test the impact of environmental differences and geographic distance on parasite genetic distance. To account for model overfitting and multicollinearity, we performed two variable selection approaches (mod-A, mod-M) (Supp. Methods). Environmental Niche Modeling (ENM) was done based on present-day and past bioclimatic variables using Maxent, as implemented in the ‘dismo’ R package ^73, 74^. A more detailed description on the landscape genomics analyses (variable selection, RDA, GDM and ENM) is presented in the Supplementary method section.

### Bioinformatics and phylogenetic analyses of LRV1

Raw sequencing reads were trimmed and filtered with fastp^75^ using the following settings: a minimum base quality (-q) of 30; the percentage of unqualified bases (-u) set to 10; per read sliding window trimming based on mean quality scores (-5, front to tail; -3, tail to front) with a window size (-W) of 1 and mean quality score (-M) of 30; right-cutting reads (-- cut_right) per 10bp windows (--cut_right_window_size) when mean quality score (-- cut_right_mean_quality) is below 30; only considering reads between 100 (-l) and 150bp (-b). LRV1 sequences were assembled *de novo* with MEGAHIT^76^ and identified using BLASTn^77^ against conventional LRV1 reference genomes LRV1-1^78^ and LRV1-4^79^ (accession numbers M92355 and U01899, respectively). In order to check and improve the quality of the assemblies, trimmed reads were mapped against the LRV1 contigs with SMALT as described above, with the minimal nucleotide identity (-y) set to 95%. Alignments were examined with Artemis^80^ and used to improve assemblies with Pilon v.1.23^81^. Genome sequences were aligned using MAFFT v.7.49 ^82^.

For comparative purposes, we included 26 (near-) complete LRV1 genomes and 13 partial LRV1 sequences of *Lb* (N=8), *Lg* (N=26) and *Lsh* (*L. shawi*) (N=1) from French Guiana, Brazil and Suriname ^43, 45, 78, 79^. Multiple sequence alignments were generated by trimming and re- aligning i) all available genomes to 5,189 bp sequences (N=57) and ii) all available sequences to 755 bp sequences (N=70). Maximum likelihood (ML) trees were generated using IQtree v.1.6.12 ^83^ with 100 bootstrap replicates and implementation of the ModelFinder function ^84^ to determine the best substitution model based on the lowest Bayesian Information Criterion (BIC). The best substitution model identified for the genome alignment was the GTR+F+R4 model, which was also applied on the partial sequence alignment for consistency. Pairwise genetic distance among LRV1 genomes were calculated with the ‘ape’ R-package ^85^ (model= ‘raw’). Viral genomes with a genetic distance below 0.09 (i.e. 9% of the sites that are dissimilar among two genomes) were grouped into distinct viral lineages. Nucleotide diversity and Fst statistics were calculated within and between viral lineages using the ‘PopGenome’ package in R^86^ while recombination was tested by pairwise homoplasy index (PHI) tests implemented in SplitsTree ^62, 87^.

### Co-phylogenetic analysis of LRV1 and *Leishmania*

Co-phylogenetic analyses were done at both the species (between LRV1 infecting *Lb* and *Lg*) and at the population (LRV1 infecting *Lb* from Peru and Bolivia) level. These analyses were performed using the phytools R package ^88^. To assess the co-evolutionary history of LRV1 with *Lb* and *Lg*, we added 24 LRV1 genomes and 19 *Lg* and 1 *Lb* SNP genotypes from French Guiana (N=19) and Brazil (N=1)^45^ (*Lg* reads accession: PRAJNA371487; LRV1 genome accessions: KY750607 to KY750630). The phylogenetic trees were tested for topological similarity calculating the Robinson-Foulds distance ^89^ between both trees with comparison against a null distribution of 1,000 permuted un-correlated trees. For LRV1 we reconstructed a ML tree as described above, including the 23 LRV1 genomes of *Lg* and 32 LRV1 genomes of *Lb*. For *Leishmania*, sequence reads of *Lg* were mapped against the M2904 reference genome and GATK Haplotypecaller was run as described above. We then performed joint genotyping on a dataset including the 19 *Lg* genomes and 80 *Lb* genomes. SNPs were filtered following similar criteria as described above (QD<2, FS>200.0, SOR>3.0, MQ<40.0, MQRankSum<-12.5, ReadPosRankSum<-8.0, QUAL<250, info DP < 10, ClusterSize=3 and ClusterWindowSize=10). The *Leishmania* ML tree was generated using IQtree based on 7,571 jointly called bi-allelic SNPs with 100 bootstrap replicates and GTR+F+R5 as best performing substitution model ^83, 84^. For the co-phylogenetic reconciliation at the intraspecific level, we focused on phylogenies generated for our dataset of LRV1 (GTR+F+R4 ML tree generated with IQtree) and *Lb* (bifurcating tree generated by fineSTRUCTURE) from Peru and Bolivia.

## DATA AVAILABILITY

Genomic sequence reads of the 79 sequenced *Lb* genomes are available on the European Nucleotide Archive (https://www.ebi.ac.uk/ena/browser/home) under accession number PRJEB4442. The assembled sequences of the 31 LRV1 genomes are available on GenBank (https://www.ncbi.nlm.nih.gov/genbank/) under accession numbers OQ673070-OQ673100.

## SUPPLEMENTARY METHODS

### Landscape genomic analyses of *Leishmania braziliensis* (*Lb*) parasites

To investigate the spatio-environmental impact on genetic variation among the three *Lb* populations, we extracted the 19 bioclimatic variables of the WorldClim2 database ^1^. All 19 variables were extracted per locality from 1 km spatial resolution raster maps after all layers were transformed to the same extent, resolution, and coordinate reference system (WGS 1984). Geographic distances among sampling points were calculated as great-circle distances using geodist R-package (measure= ‘haversine’ ^2^).

The impact of geographic distance on the genetic differentiation of *Lb* populations was assessed through Mantel tests between i) interdeme geographic distance and the Weir-Cockerham’s F_ST_^3^; ii) the geographic distance and Bray-Curtis genetic dissimilarity among individuals. The environmental and geographic influence on the ancestral genomic population structure of *Lb* was disentangled through redundancy analysis (RDA) and generalized dissimilarity modeling (GDM) including geographic distance and a selection of bioclimatic variables to reduce model overfitting and multicollinearity. Variable selection was performed adopting two approaches: i) mod-A: an RDA-based forward selection procedure using ‘ordiR2step’ function of the vegan R-package ^4, 5^; ii) mod-M: a manual variable selection procedure, selecting variables if their added contribution increased the adjusted R-squared and if both the overall RDA model and individual variable were significant (p-val < 0.05).

Variation partitioning through RDA analysis was performed on Hellinger transformed SNP data, longitude, latitude and standardized environmental variables, using the ‘decostand’ and ‘rda’ functions from the ‘vegan’ R-package^4^, to disentangle the influence of geography (isolation-by-distance) and climate (Isolation-by-environment). Finally, the different variance components were compared based on their adjusted R-squared and each explanatory component was tested for significance using the vegan::anova.cca function. In addition to RDA variation partitioning, we constructed generalized dissimilarity models (GDMs) using the gdm R-package^6^, to investigate spatio-environmental patterns of the *Lb* genetic variability in a non-linear way ^6, 7^. The GDMs constituted a genetic distance matrix (Bray-Curtis dissimilarity of Hellinger transformed SNP genotypes) as response variable and the bioclimatic variables (as selected by the mod-A and mod-M variable selection procedures) as explanatory variables. We accounted for geographic distance effects on the genomic variability by fitting two GDMs per variable selection approach including and excluding the inter-individual geographic distance matrix. Relative variable importance in each GDM was estimated based on the I-spline basis function (i.e., the maximum height of the response curves) along with uncertainty assessment by performing 1000 iterations of each GDM model ^8^.

Based on the most important environmental variables influencing the *Lb* population genomic structure, we attempted to estimate and map patches of suitable habitat for *Lb* within our study region based on present-day^1^, LGM^9, 10^ and LIG^10, 11^ bioclimatic data. Ecological niche models (ENMs) were constructed using Maxent, as implemented in the dismo R-package ^12, 13^ for both environmental variable selection methods with the following parameters: linear, quadratic, product, threshold, hinge, 10 cross-validation replications and regularization multiplier (rm) set to 1, 1.5 and 2. Habitat suitability maps were constructed by averaging the predictions from all 10 replicates on present-day, LGM or LIG environmental data. A jackknife procedure was included to measure relative variable contribution and importance.

## SUPPLEMENTARY RESULTS

### Landscape genomics of *Lb* parasites

Upon the strong signatures of geographical isolation of the three ancestral *Lb* components (Fig 2A) and the lack of association between the inter-population (great-circle) geographic distance and the Weir & Cockerham’s *F*_ST_ (Supp Fig. 18; Supp Table 13), we investigated the differential influence of geography and environmental variables on the *Lb* population structure in the region. When addressing the inter-individual association of geographic distance with genetic distance (Bray-Curtis dissimilarity of SNP genotypes) we picked up a pattern resembling case-IV isolation-by-distance (i.e., increasing genetic distance with geographic distance up to a certain point after which the relationship weakens down; Supp. Fig. 19; Supp. Table 14). This revealed that isolation-by-distance mainly plays a role within populations over distances up to ca. 500km, while IBD diminishes on an inter-population level when geographic distances become too great (> 500km).

To investigate what other factors besides geography influenced the population divergence among the *Lb* populations, we investigated the potential impact of the abiotic environment through RDA-based variation partitioning and GDM analysis including geographic distance and 19 bioclimatic variables (Supp. Table 6). Two variable selection approaches were adopted to reduce model overfitting and multicollinearity among bioclimatic variables (Supp. methods). The mod-A variable selection initially resulted in six variables, although they revealed large variance inflation factors (vif) which prompted us to remove variables with a vif > 10 in a stepwise manner, retaining only two variables: ‘isothermality’ (bio3) and ‘Precipitation of the driest month’ (bio14). In contrast, the mod-M approach resulted in a final selection of five variables, each with vif < 10: ‘isothermality’ (bio3), ‘Precipitation of the driest month’ (bio14), ‘precipitation of warmest quarter’ (bio18), ‘precipitation seasonality’ (bio15), ‘Annual mean diurnal range’ (bio2). (Supp. Table 7).

Variation partitioning of the automated variable selection model revealed that about one-third (27.3%) of the total genomic variability could be explained by the environment (bio3, bio14) and geography together, of which both components contributed 10.2% and 7.5%, respectively (Supp. Fig 8A; Supp. Table 8). In addition, the remaining 9.6% of the explained genomic variability indicates a strong confounding effect among the environment and geography components with the RDA model, meaning that about one-third of the explainable genomic variation cannot be attributed to one specific explanatory component (Supp. Table 8). In parallel, generalized dissimilarity models (GDM), including the same variables, revealed similar patterns in explaining the genomic variability by the environment (bio3, bio14) relative to geography (Supp. Fig 20). Here, GDMs could explain 55.86% (excluding geography) to 65.38% (including geography) of the genomic deviance (null deviance).

In contrast, variation partitioning of the RDA-model based on the manual variable selection approach (bio2, bio3, bio14, bio15, bio18) revealed a stronger environmental contribution in explaining the genomic variation in *Lb*. From the full RDA-model, explaining 34.9% of the total genomic variability, about 51% could be explained by the entire environmental component whereas only 5.4% could be explained by geography (Supp. Fig 8B; Supp. Table 8). Additional GDM, explaining 59.98% (excluding geography) to 65.04% (including geography) of the genomic deviance, gave consistently similar results to the RDA model (Supp. Fig 20).

In accordance with the different variation partitioning models (RDA and GDM), revealing the key role in explaining the *Lb* population divergence, present-day habitat suitability predictions showed that regions of suitable (abiotic) habitat for *Lb* coincided with tropical rainforests, as predicted by the Koppen-Geiger climate classification, where the three ancestral *Lb* populations were surrounded by less suitable tropical monsoon forests (Fig 2B,C). Additional suitability predictions using Last Glacial Maximum (LGM) and Last Interglacial (LIG) periods revealed similar regions of suitable habitat, suggesting the suitable regions for ancestral *Lb* populations have been relatively stable over the past 120,000 years (Supp. Fig 9, 10).

### Quality assessment of the LRV1 genome assemblies

The LRV1 genomes included in this study were generated through dsRNA extraction from 31 LRV1-positive *L. braziliensis* isolates that were re-cultured following total RNA sequencing, *de novo* assembly and LRV1 contig extraction by a local BLAST search (see methods). This procedure failed for two *Lb* isolates, either due to difficulties during culturing (PER096) or because the assembly yielded a partial LRV1 genome (PER231). Two *Lb* isolates (CUM65 and LC2321) each harbored two LRV1 genomes, differing at 999 (for CUM65) and 60 (for LC2321) nucleotides, bringing the total to 31 viral genomes. The assembly quality of the genomes was examined by investigating the coverage of mapped (paired) reads, SNP counts after mapping, comparison with analogous genomic regions to ∼1kb sequences obtained through conventional Sanger sequencing and by read-based computational assembly improvement using Pilon ^14^.

From the average 0.04% of reads that mapped against their respective LRV1 contig (Supp. Table 9), an average of 85.7% (81.3% - 91.5%) of the reads were properly paired (i.e., correctly oriented reads with respect to each other and with proper insert sizes). In addition, most LRV1 strains contained few SNPs (zero to three) of which all were heterozygous except one. PER130 showed one homozygous SNP, which was located at the 5’ end (position 6) of the assembled genome which was trimmed off in downstream analyses. However, in LRV-Lb- LC2321 we encountered 51 heterozygous SNPs (data not shown), suggesting the possibility of a mixed viral infection (i.e. two LRV1 strains present in one parasite isolate of which one is much lower in abundance than the other). To examine this in more detail, we extracted the reads mapping to the potential chimeric LRV1 genome and re-assembled the reads using a recently developed strain-resolving *de novo* assembler ^15^, developed to extract various viral strains from mixed infection samples. This resulted in two LRV1 contigs: LC2321.1 (5,260bp) and LC2321.2 (4,738bp), and considerably dropped the number of heterozygous SNPs found in both strains. LC2321.1 showed nine heterozygous SNP of which eight were located on a non-resolved part of the LC2321.2 genome. LC2321.2 on the other hand did not reveal any SNPs. Furthermore, the remaining quality statistics of both strain resolved genomes from LC2321 were similar to the other assemblies (Supp. Table 9).

The sequence identity of the assembled genomes was assessed by comparison with analogous ∼1kb (1197bp) sequences obtained through conventional Sanger Sequencing. This revealed for 83.8% (26/31) of the genomes a sequence identity of 100% (Supp. Table 9). The remaining five genomes encompassed both genomes of CUM65 and LC2321 (mixed LRV1 infections) and PER212. For CUM65, we observed a sequence identity of 99% (12 mismatches) and 86% (152 mismatches) for LRV1-Lb-CUM65.1 and LRV1-Lb-CUM65.2, respectively. For LC2321, both resolved strains showed a sequence identity of 99% with three and ten nucleotide mismatches in LRV1-Lb-LC2321.1 and LRV1-Lb-LC2321.2, respectively. These mismatches were unique to each strain, which might indicate the Sanger sequence is a chimera of both strains. Finally, the 99% identity of PER212 with its respective partial Sanger sequence showed only one nucleotide mismatch, not corresponding to the identified heterozygous SNP, suggesting a badly called base during the Sanger sequencing.

---

**Supplementary Figure 1.**
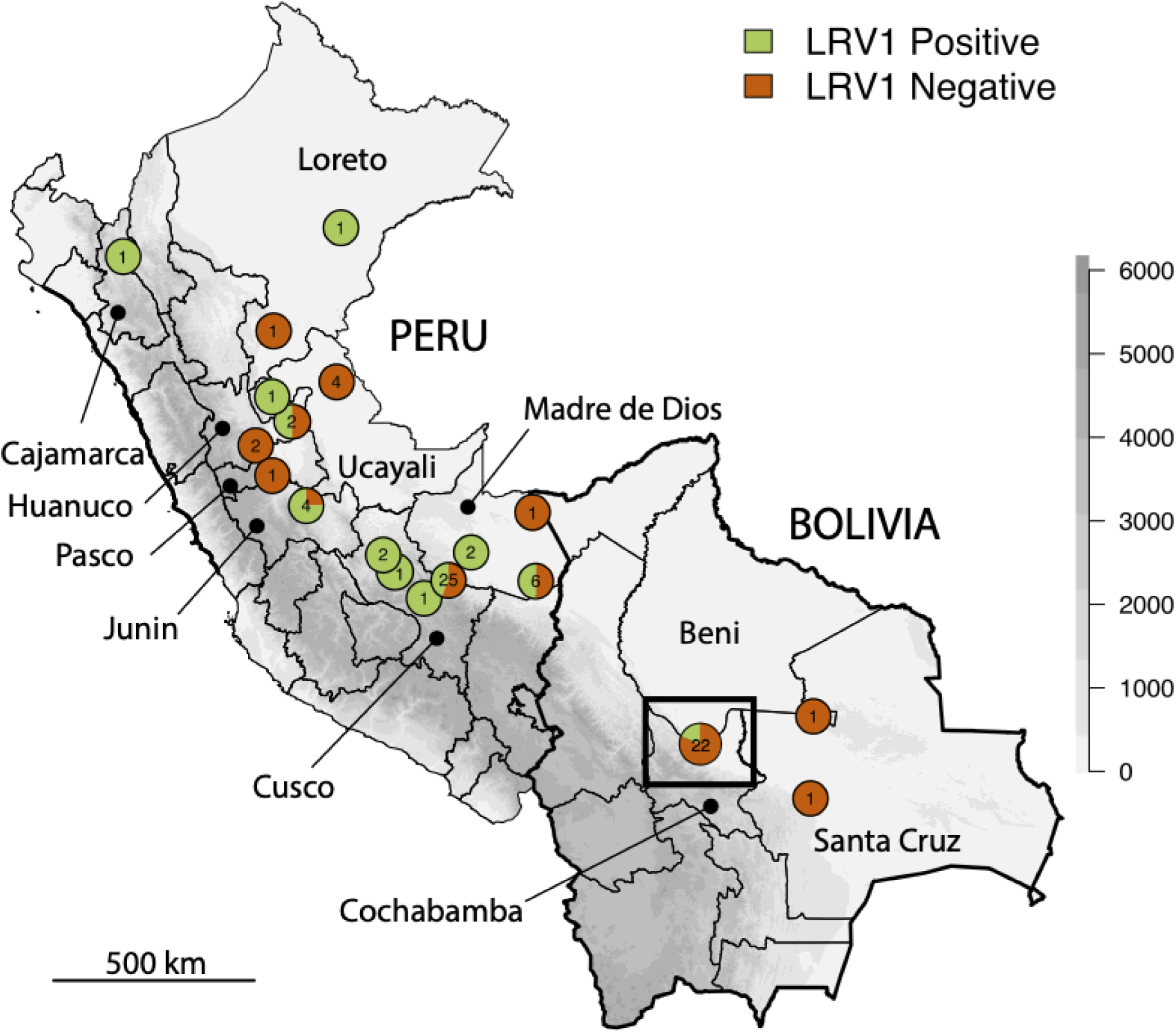
Geographic origin of 79 *Lb* isolates from Peru and Bolivia that were included in this study. Rectangular box indicates the location of the Isiboro National Park that extends between the Departments of Cochabamba and Beni. Gray-scale represents altitude in meters.

**Supplementary Figure 2.**
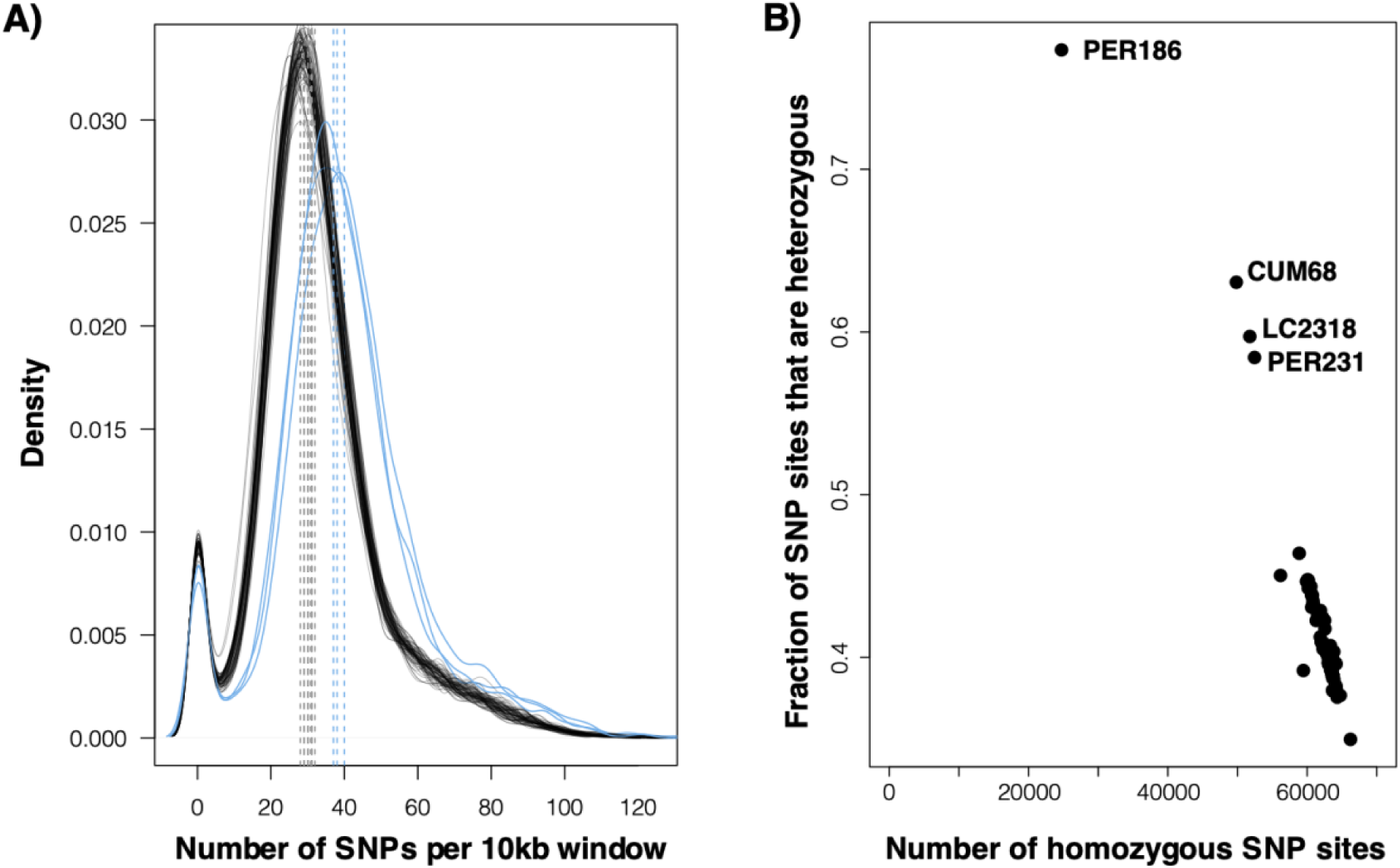
**A)** Kernel density plots of the number of SNPs per 10kb window for each of the 79 *Lb* genomes. The median number of SNPs per 10kb window is indicated with gray vertical dashed lines and ranges between 28 and 32 SNPs for the majority of isolates. Three isolates showed slightly larger SNP densities (indicated with blue lines in the plot): 37 SNPs in PER231, 38 SNPs in LC2318 and 40 SNPs in CUM68. **B)** Fraction of SNP sites that are heterozygous versus the number of homozygous SNP sites in each of the 79 *Lb* genomes.

**Supplementary Figure 3.**
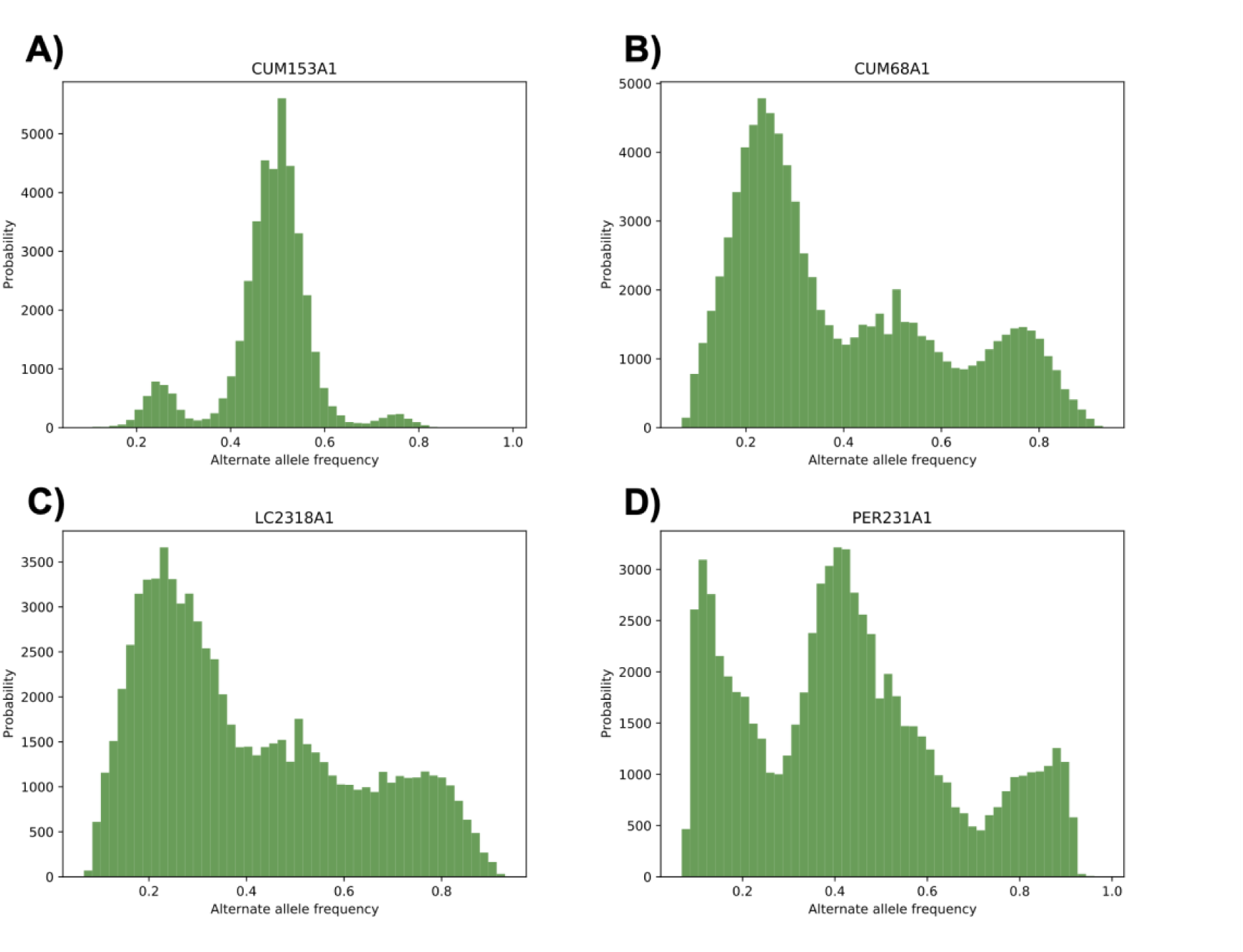
Genome-wide distribution of alternate allele read depth frequencies at heterozygous sites. **(A)** Example of a largely diploid individual (CUM153) with allele frequencies centered around 0.5, which was observed for 77/79 *Lb* genomes included in this study. **(B-C)** Two isolates (CUM68 and LC2318) were symptomatic of tetraploidy, with modes of allele frequencies equal to 0.25, 0.5 and 0.75. **(C)** One isolate (PER231) showed a skewed distribution, suggesting that it may be the result of a mixed infection or contamination.

**Supplementary Figure 4.**
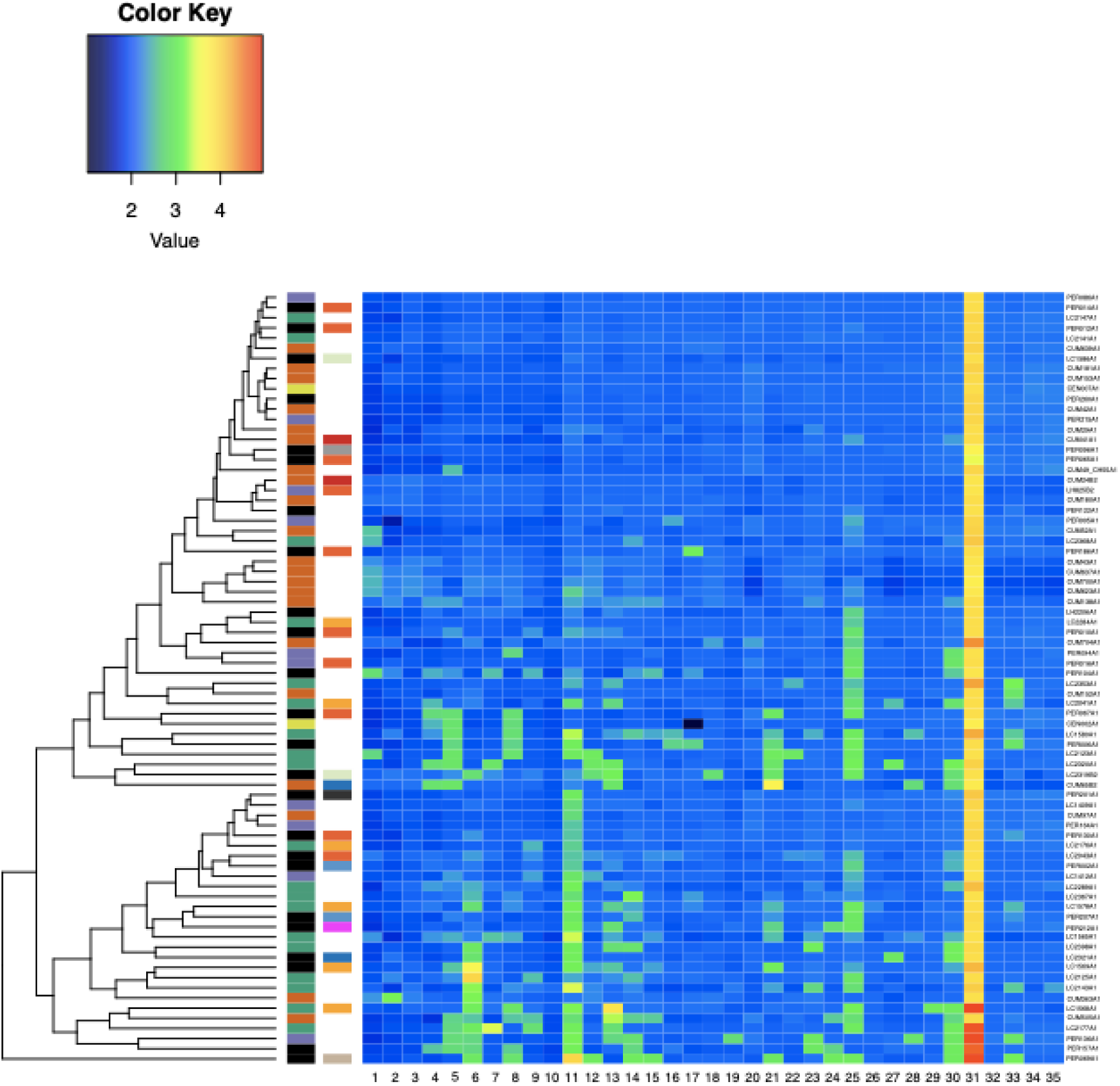
Variation in chromosome copy numbers in a panel of 76 *Lb* genomes. Isolates are clustered according to similarity in somy estimation with aneuploid individuals at the bottom of the heatmap and overall diploid individuals at the top of the heatmap. Coloured boxes on the left of the heatmap represent: *Left -* the inferred *Lb* populations PAU (green), INP (orange), HUP (purple), STC (yellow-green) and ADM (black); *Right* - the identified LRV1 lineages encountered in each *Lb* isolate. I (orange), II (dark gray), III (pink), IV (steelblue), V (yellow), VI (beige), VII (dark blue), VIII (red), IX (light green), NA (light gray).

**Supplementary Figure 5.**
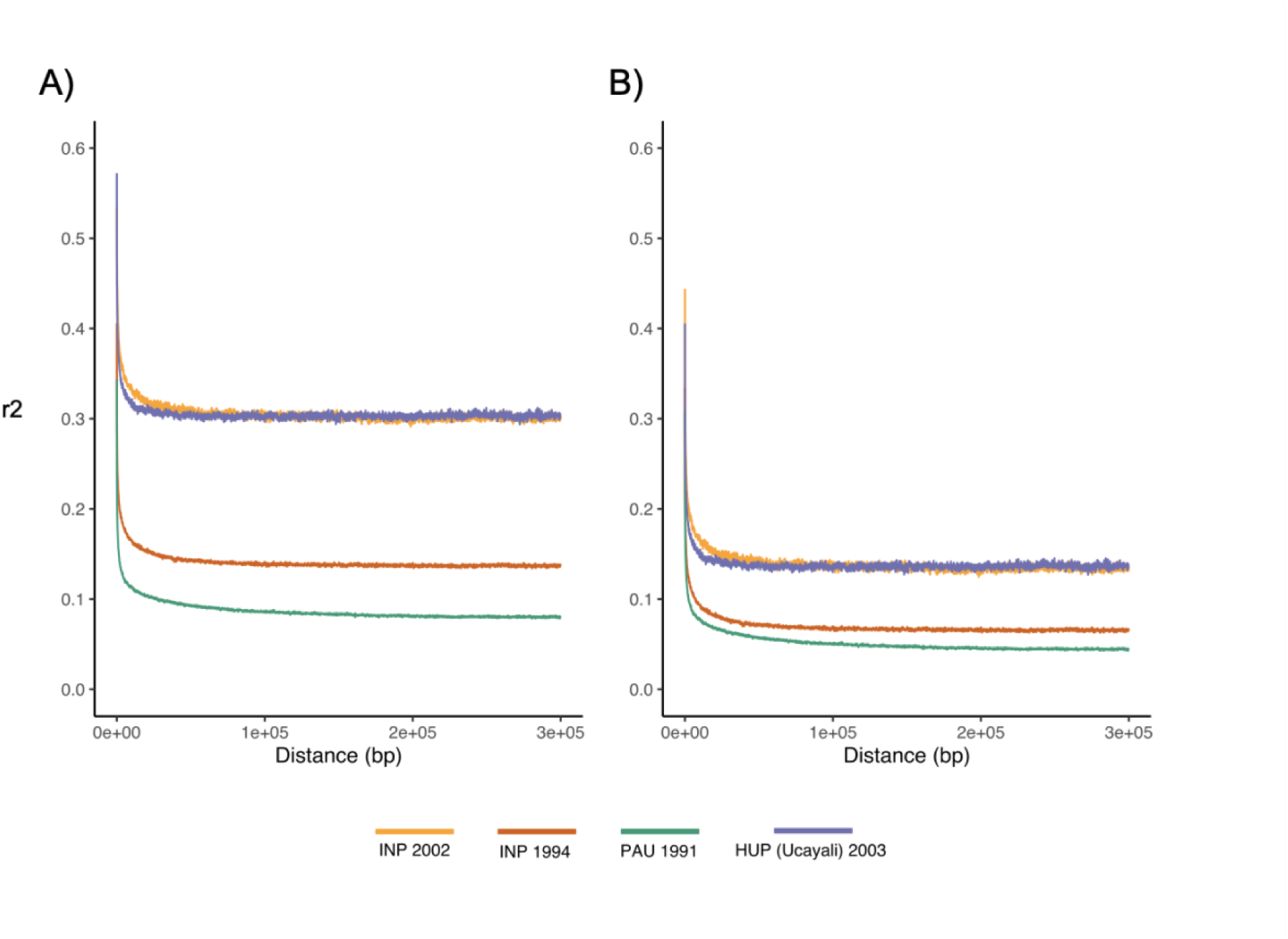
Linkage Disequilibrium decay plots after correction for population structure and spatio-temporal Wahlund effects. **A)** Uncorrected for sample size. **B)** Corrected for sample size (r2 - 1/(2n)) ^16^.

**Supplementary Figure 6.**
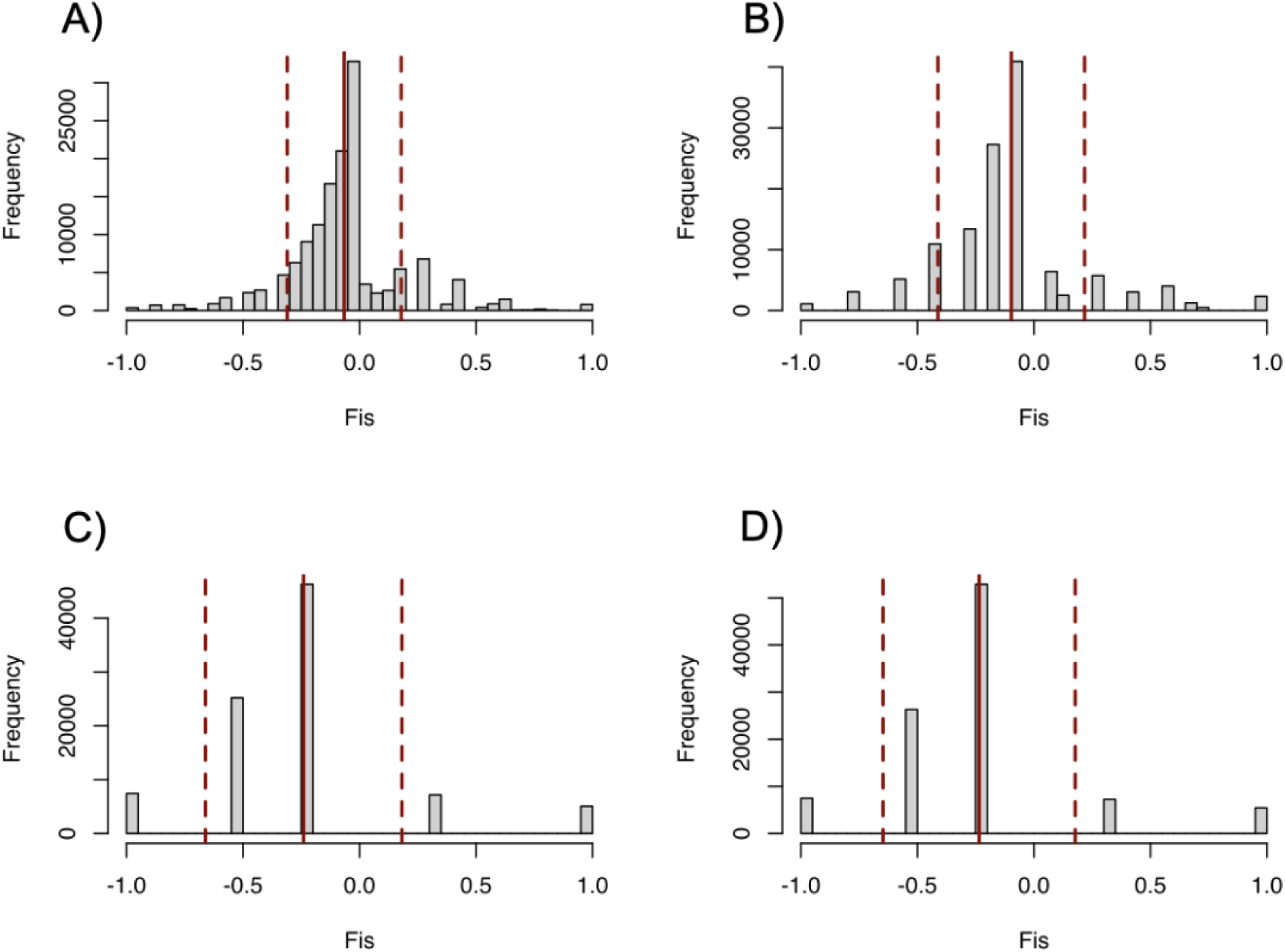
Fis distributions after correction for population structure and spatio- temporal Wahlund effects. **A)** Individuals from PAU sampled in 1991 (N=14). **B)** Individuals from INP sampled in 1994 (N=7). **C)** Individuals from INP sampled in 2002 (N=3). **D)** Individuals from HUP (Ucayali) sampled in 2003 (N=3). Solid red lines depict the mean Fis value for each population. Dashed red lines represent the mean’s standard deviation.

**Supplementary Figure 7.**
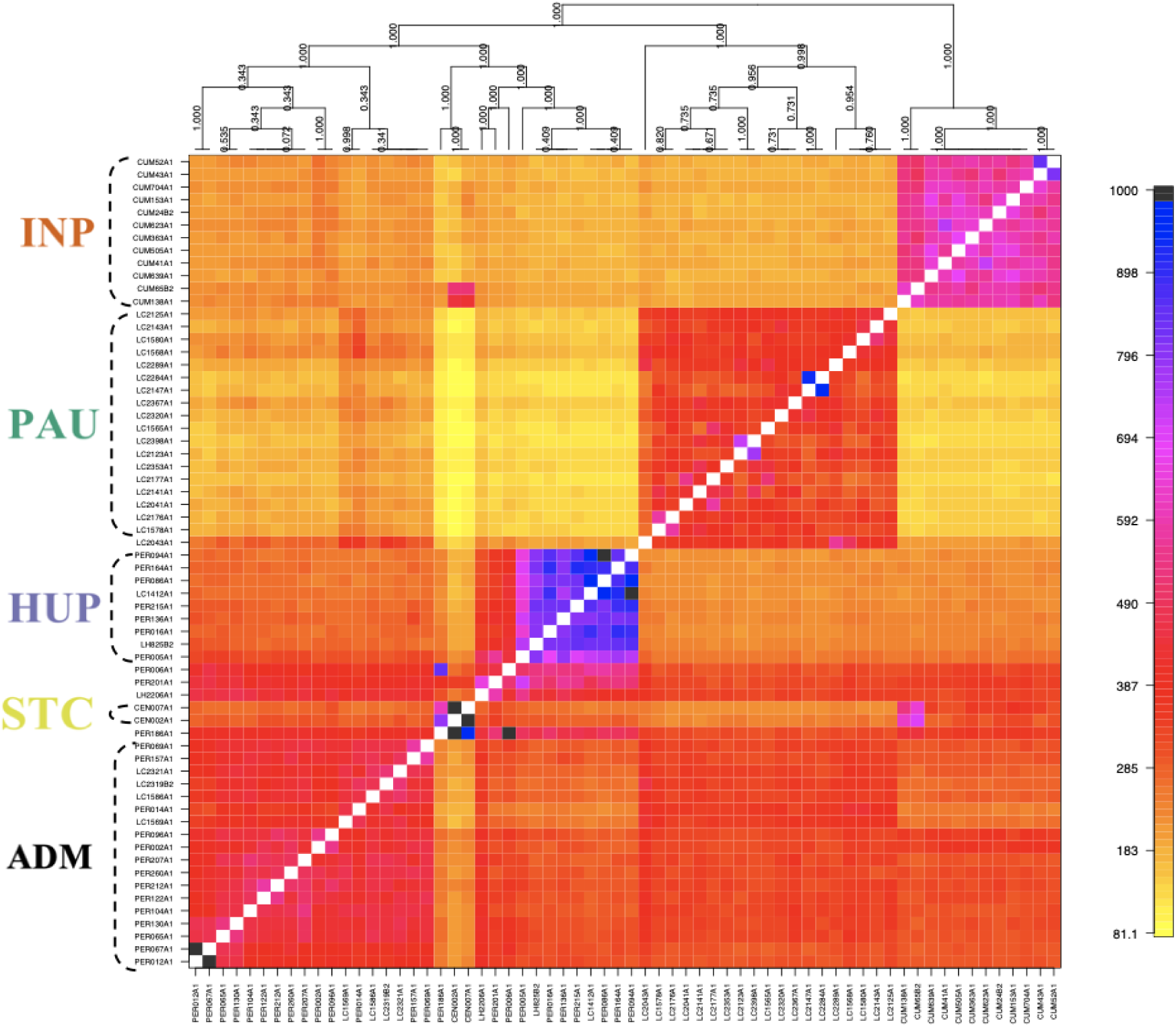
Co-ancestry matrix, as inferred by fineSTRUCTURE, depicting the pairwise number of received (rows) and donated (columns) haplotype segments between two parasite genomes. Color key on the right shows the amount of shared haplotype segments and is capped on 1000 for visibility reasons. Individuals are ordered according to the fineSTRCUTURE clustering outcome (above matrix) and dashed accolades indicate the three main parasite groups (INP, HUP, PAU) and the two main groups of admixed parasites (ADM, STC); the remainder of the parasites were of uncertain ancestry (UNC).

**Supplementary Figure 8:**
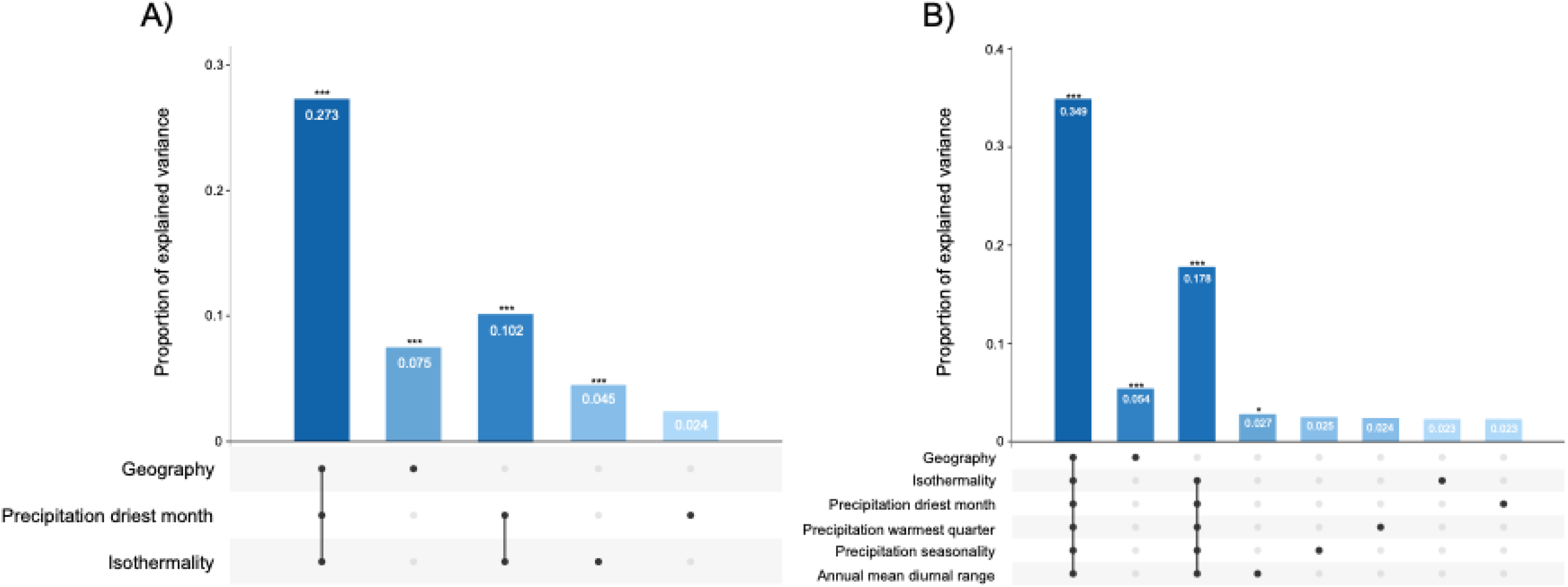
Partial RDA models showing the influence of the environment (bioclimatic variables) and geography on the genomic variability between the three inferred populations in Peru and Bolivia. A) Partial RDA model including geography, isothermality (bio3) and precipitation of driest month (bio14). Bioclimatic variables were selected based on the automated variable selection approach (suppl. Table 6; see methods). B) Partial RDA model including geography, Isothermality (bio3), precipitation driest month (bio14), precipitation warmest quarter (bio18), precipitation seasonality (bio15) and annual mean diurnal range (bio2). Bioclimatic variables were selected based on the manual variable selection approach (suppl. 6; see methods).

**Supplementary Figure 9.**
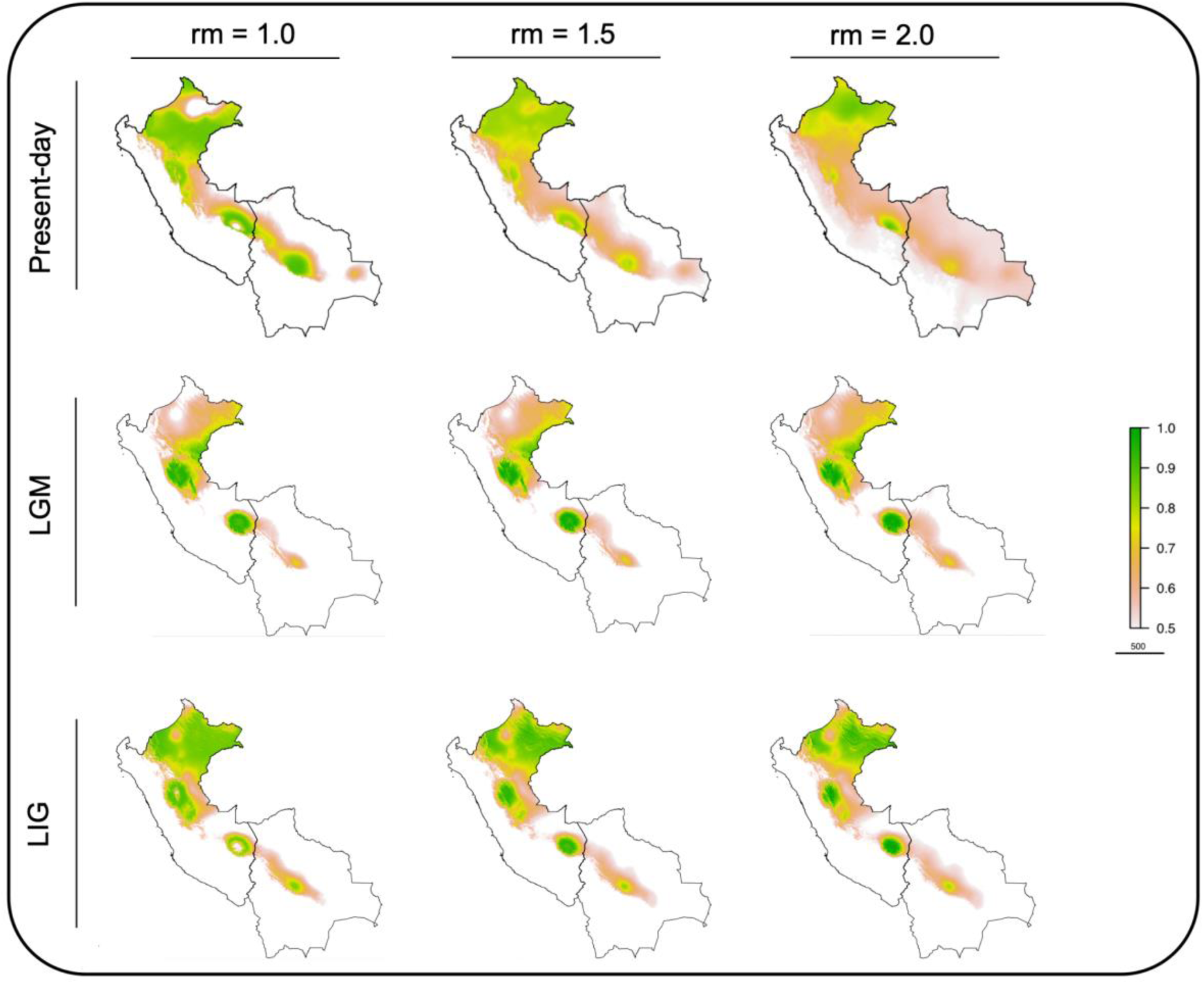
Ecological niche models based on Present-day, LGM and LIG data of isothermality (bio3) and Precipitation of the driest month (bio14) (mod-A variable selection approach) with different regularization values (rm = 1, 1.5 and 2). The continuous-scale legend represents habitat suitability (probability of occurrence).

**Supplementary Figure 10.**
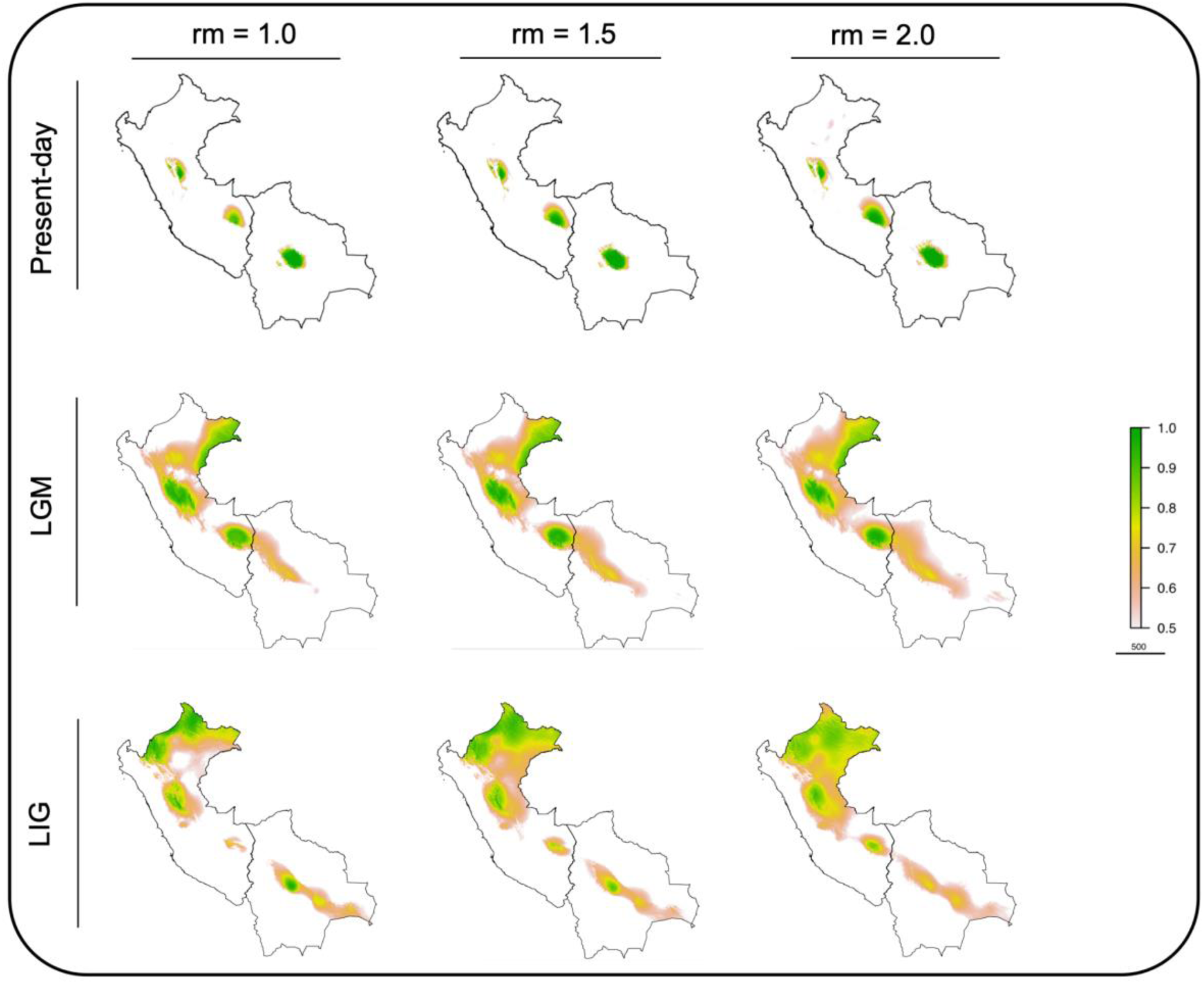
Ecological niche models based on Present-day, LGM and LIG data of isothermality (bio3), Precipitation of the driest month (bio14), precipitation of the warmest quarter (bio18), precipitation seasonality (bio15) and annual mean diurnal range (bio2) (mod- M variable selection approach) with different regularization values (rm = 1, 1.5 and 2). The continuous-scale legend represents habitat suitability (probability of occurrence).

**Supplementary Figure 11.**
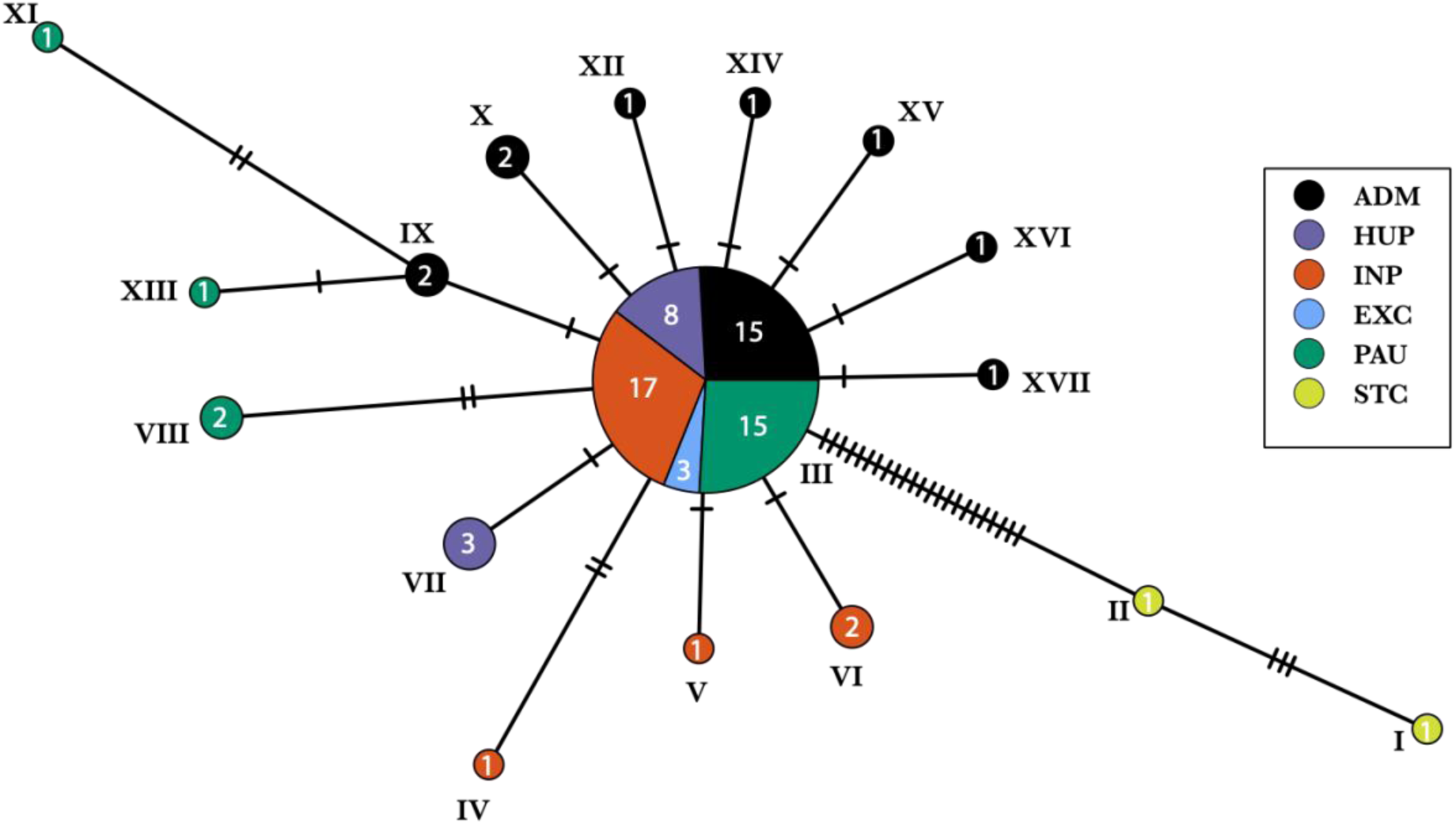
TCS Haplotype network with PopART based on 53 high-quality SNPs identified within the coding region of the haploid mitochondrial maxicircle. A total of 17 haplotypes (shown with Roman numbers) were identified within our set of 80 *Lb* isolates from Peru and Bolivia. The size of the circles represent the number of sequences that represent a given haplotype; this number is also written in white within each circle. Colors indicate the five groups of parasites as identified with ADMIXTURE and fineSTRUCTURE using genome-wide SNPs; the EXC group indicates the three isolates that were excluded from population structure analyses. The dominant haplotype III is represented by 58 maxicircle sequences (72.5% of the 80 included sequences) and is found in four of the five groups. Black bars represent the number of mutations between two haplotypes.

**Supplementary Figure 12.**
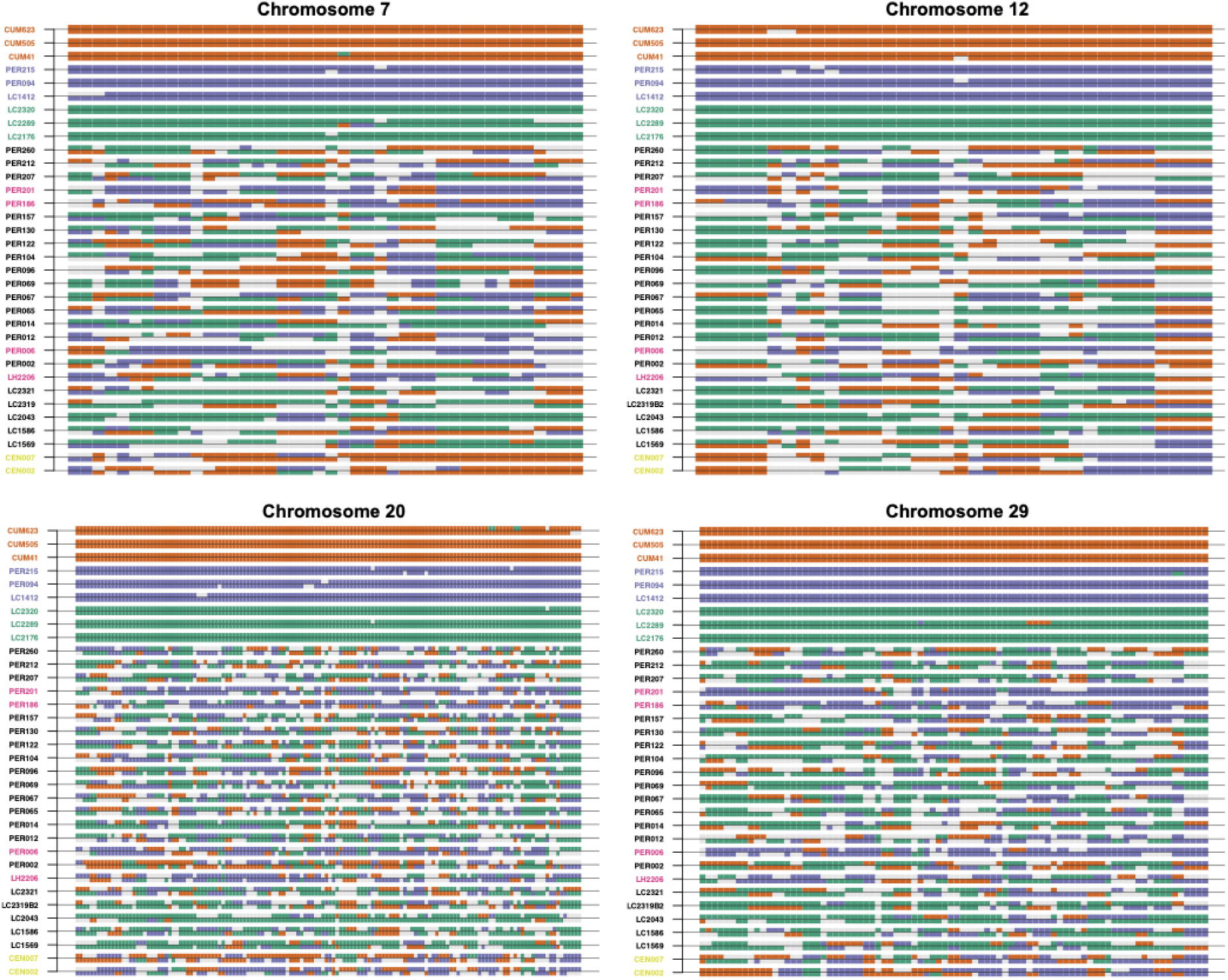
PCAdmix local ancestry assignment to PAU, INP and HUP source populations of the 19 ADM isolates, 4 UNC isolates, 2 STC isolates and three randomly selected isolates from each source population. Ancestry was assigned in windows of 30 SNPs along each chromosome (here only chromosomes 7, 12, 20 and 29 are shown).

**Supplementary Figure 13.**
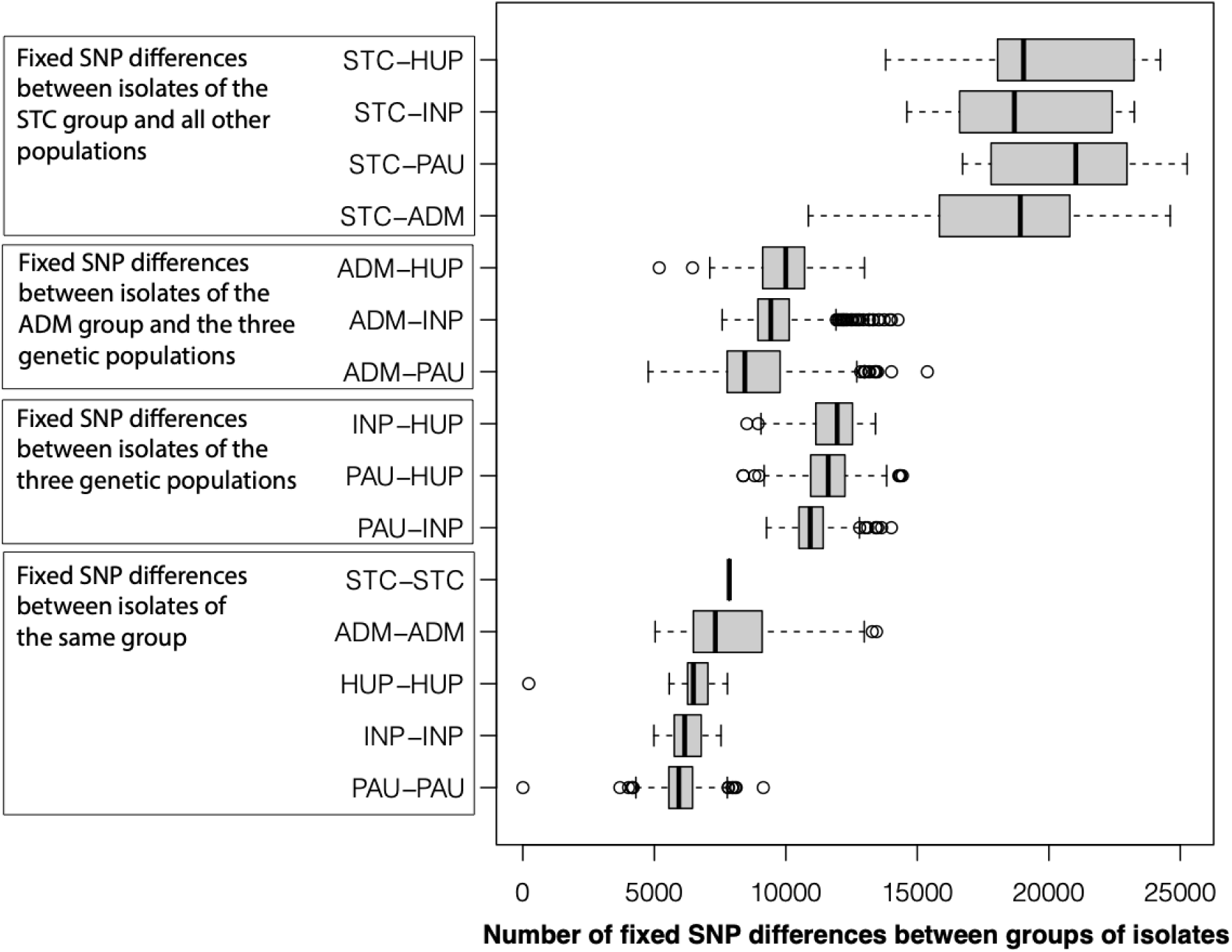
Number of fixed SNP differences between *Lb* isolates of the same group or between *Lb* isolates of different groups.

**Supplementary Figure 14.**
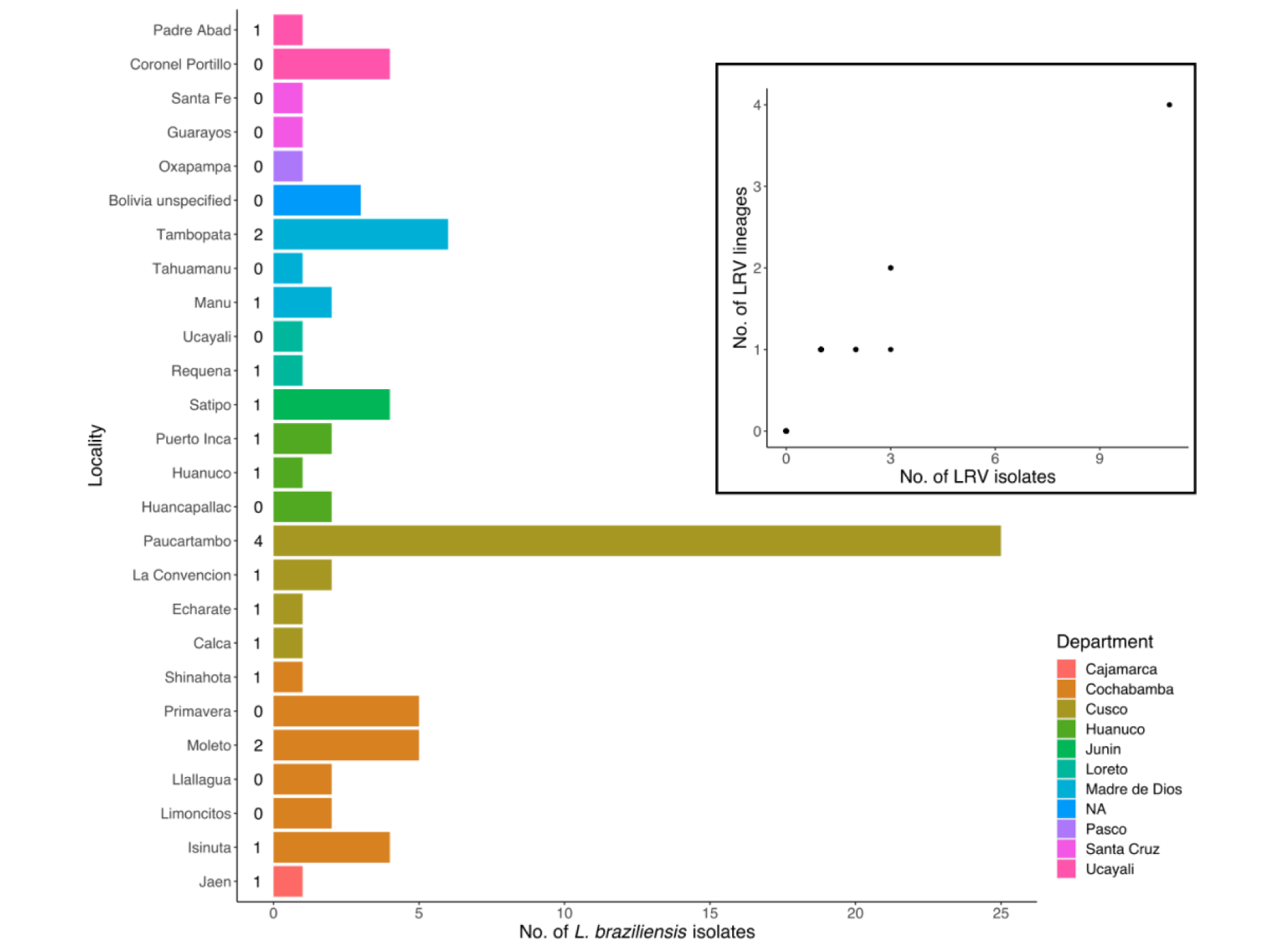
Barplot shows the number of *Lb* isolates per sampling locality. Number on the right of each bar shows the number of *Lb* isolates that were positive for LRV1. Bars are coloured according to the Department. Inset reveals the number of LRV1 lineages versus the number of LRV1 isolates that were recovered in a given locality.

**Supplementary Figure 15.**
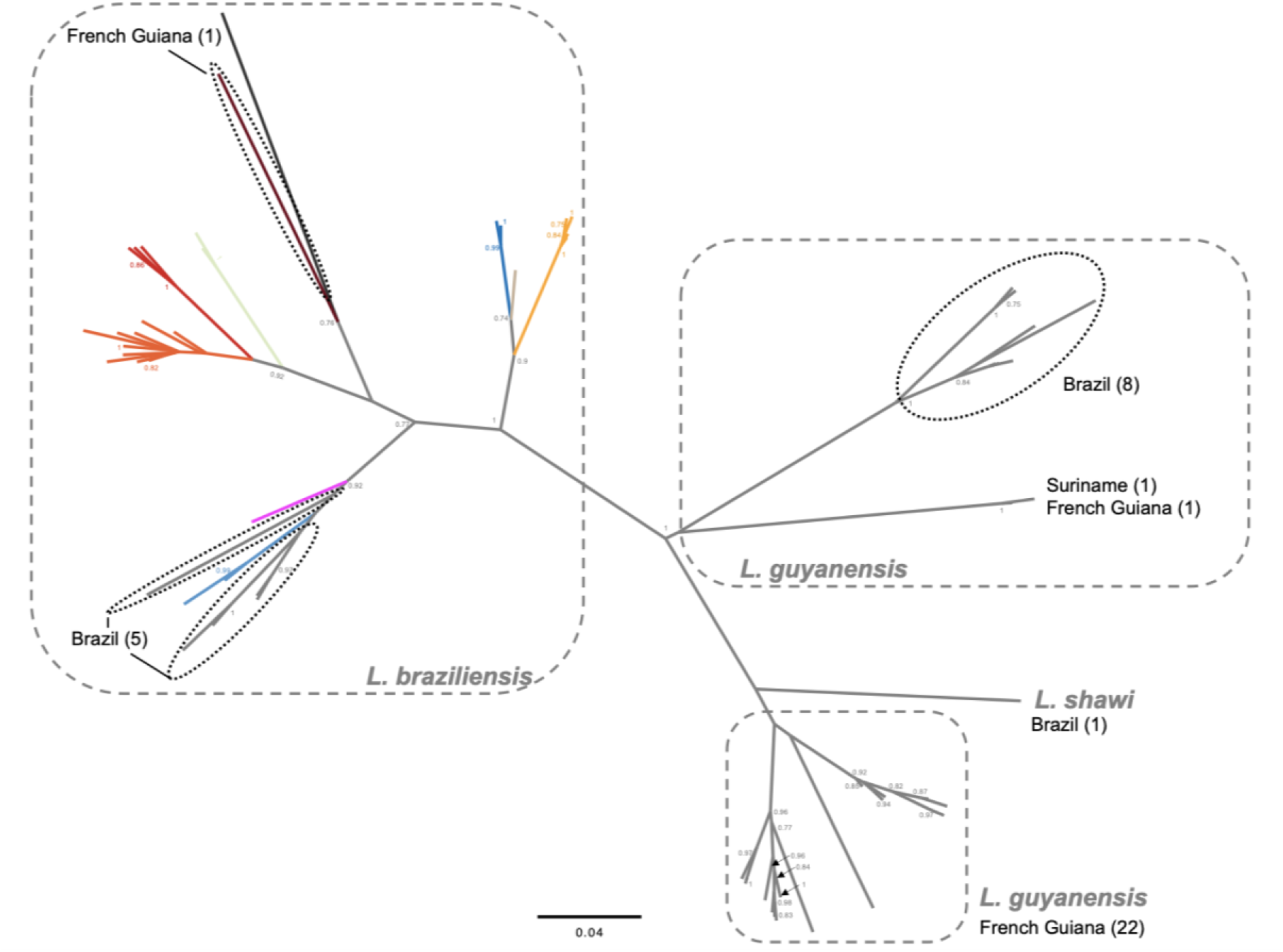
Maximum likelihood tree based on partial LRV1 sequences (756bp) from *L. braziliensis*, *L. guyanensis* and *L. shawi* originating from Peru, Bolivia, Brazil, French Guiana and Suriname ^17–20^. Colored lineages in *L. braziliensis* correspond to the LRV1 lineages described in this study (Fig. 3).

**Supplementary Figure 16.**
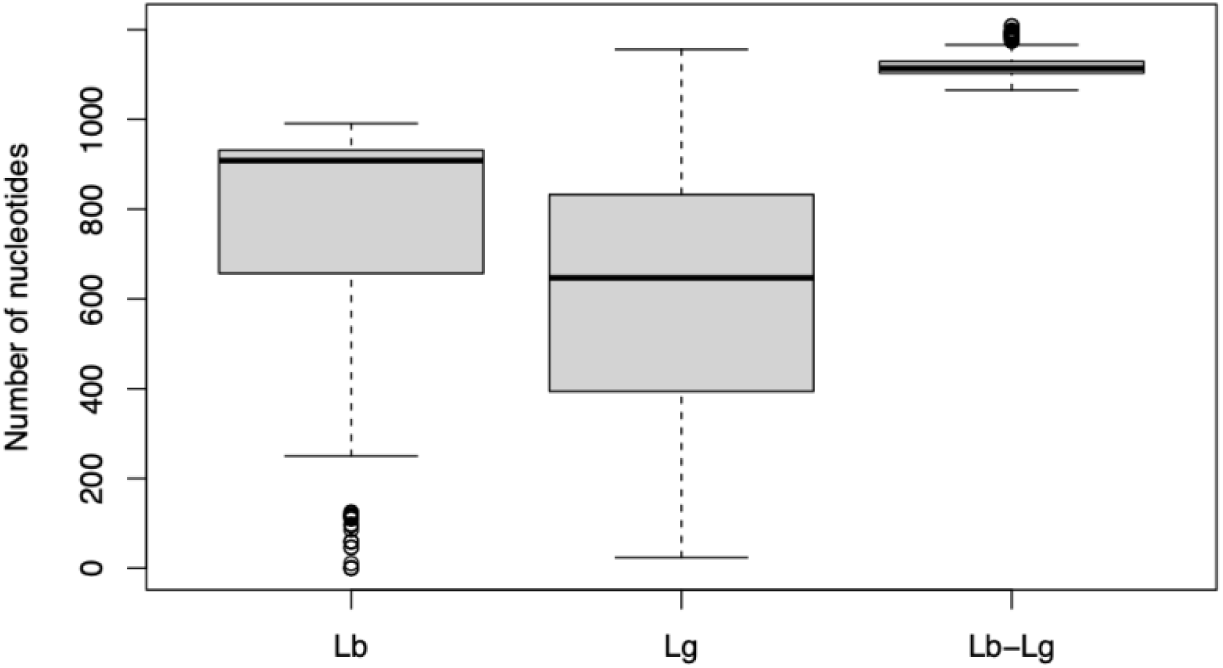
Nucleotide differences - on average higher between *Lb* and *Lg* viral genomes then within *Lb* and *Lg*.

**Supplementary Figure 17.**
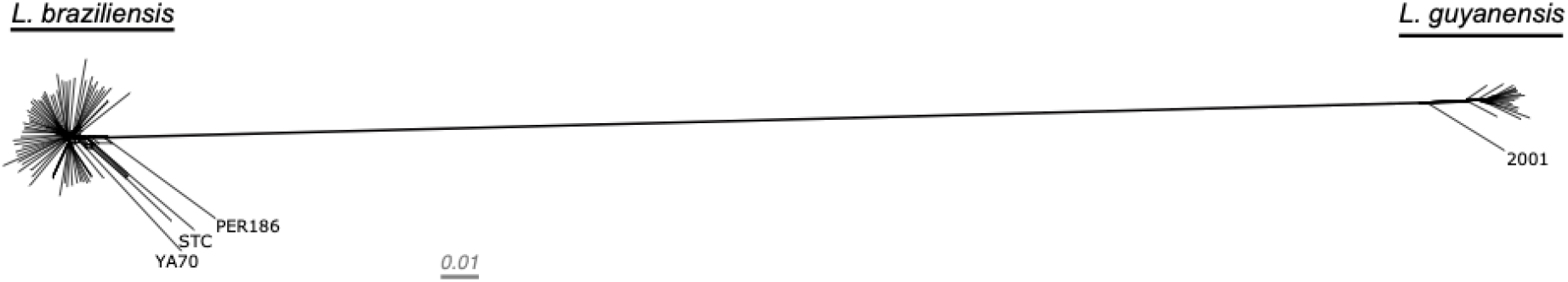
A phylogenetic network, inferred with SPLITSTREE, based on uncorrected p-distances between 77 *Lb* and 19 *Lg* isolates typed at 7,571 bi-allelic SNPs.

**Supplementary Figure 18.**
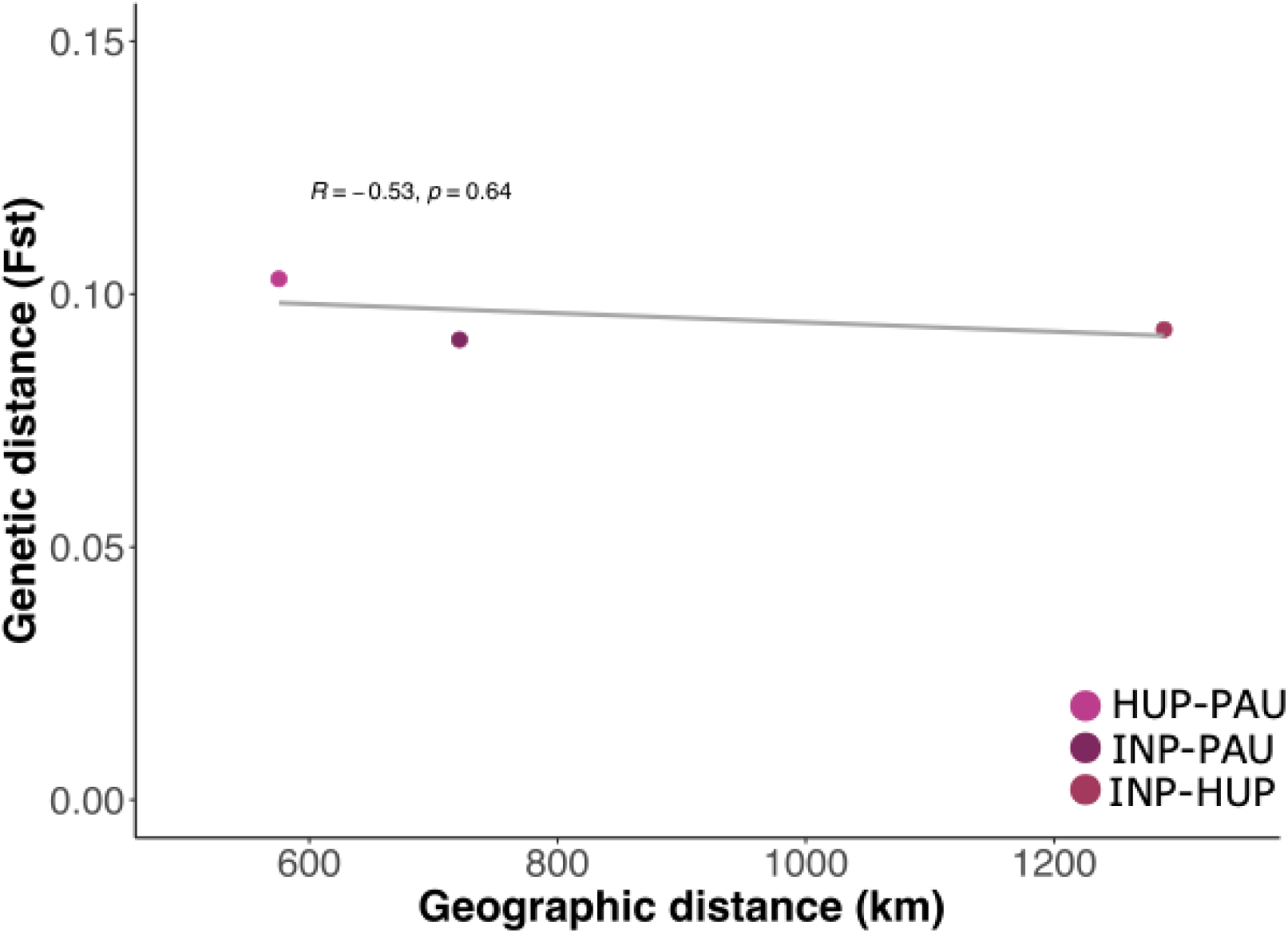
Linear regression of the interdeme great-circle distance (km) and the Weir & Cockerham’s Fst of the three ancestral *L. braziliensis* components.

**Supplementary Figure 19.**
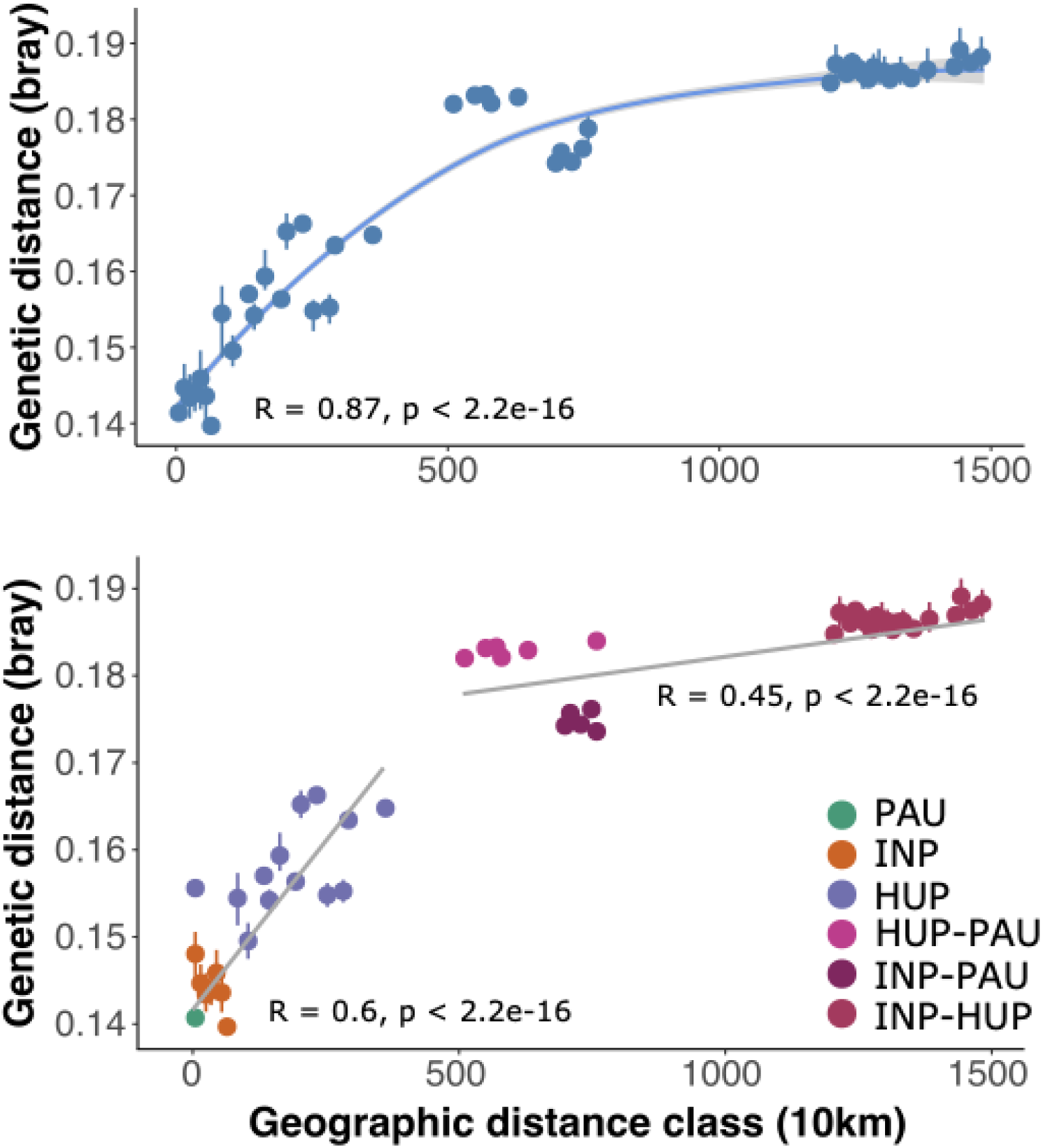
Regressions of the inter-individual great-circle distance (km) and pairwise genetic distance (Bray-Curtis dissimilarity of SNP genotypes) revealing a case-IV IBD pattern. a) Loess regression for all individuals with a loess value of 1.2. b) Linear regression of intra- and inter-population genetic distance vs. geographic distance, separately.

**Supplementary Figure 20.**
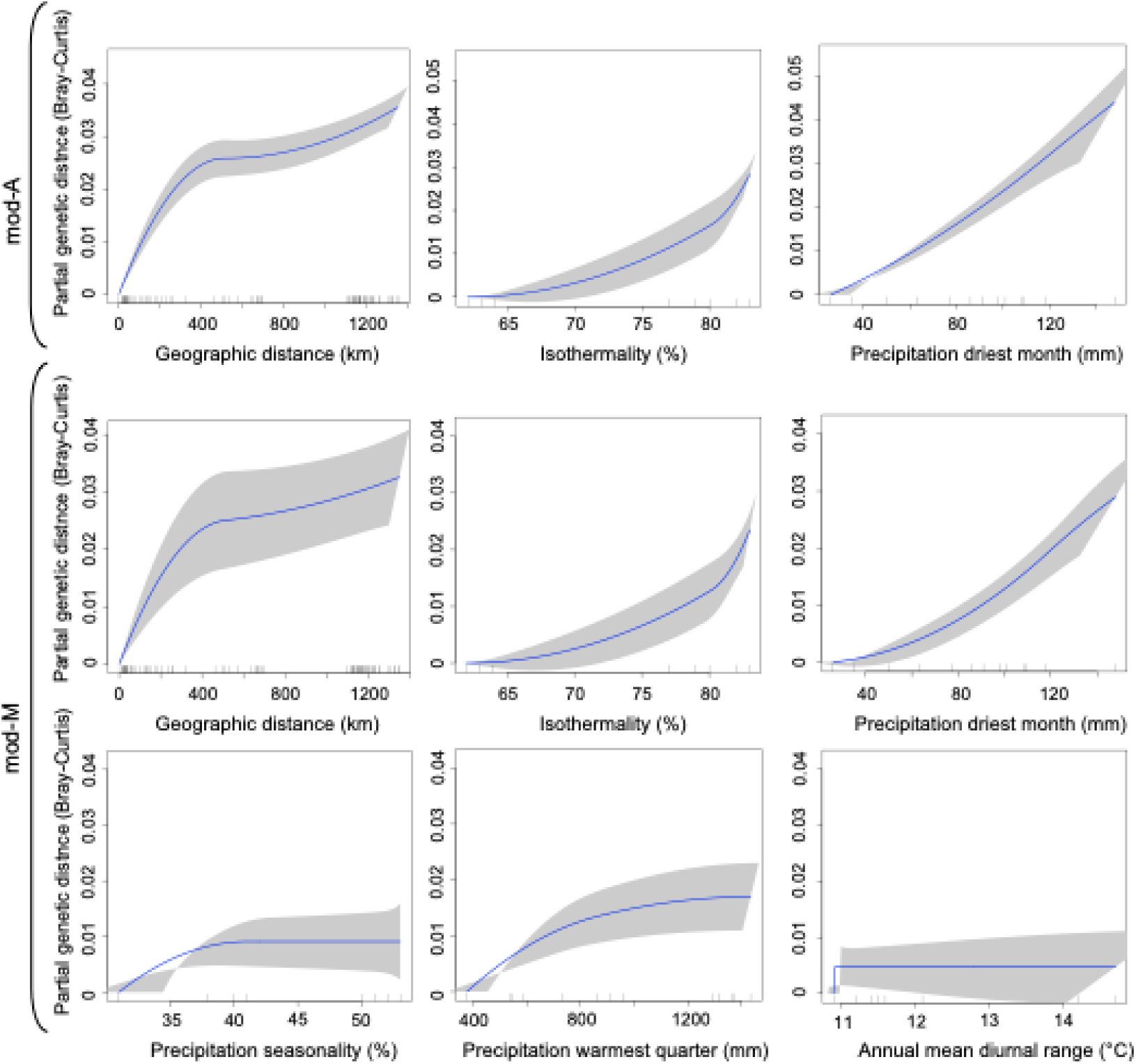
I-spline response curves for each variable included in the different generalized dissimilarity models (mod-A & mod-M). The maximum curve height represents the amount of genetic variability the variable explains (i.e.the variable importance). The curves’ slope indicates the degree of the explained genomic dissimilarity along the spatial or environmental gradient, meaning a steeper slope represents greater dissimilarity between two points while a shallower slope suggests less variability among the two points.

